# Encapsidic production and isolation of degradation-prone polypeptides

**DOI:** 10.1101/2025.02.04.636450

**Authors:** Agnieszka Gawin, Jędrzej Pankowski, Maria Zarechyntsava, Dominika Kwasna, Damian Kloska, Lukasz Koziej, Sebastian Glatt, Neli Kachamakova-Trojanowska, Yusuke Azuma

**Affiliations:** Malopolska Centre of Biotechnology, Jagiellonian University, Gronostajowa 7A, 30-387 Krakow, Poland; Faculty of Biochemistry, Biophysics, and Biotechnology, Jagiellonian University, Gronostajowa 7, 30-387 Krakow, Poland; Department for Biological Sciences and Pathobiology, University of Veterinary Medicine Vienna, 1210 Vienna, Austria

## Abstract

Degradation during production and delivery is a significant bottleneck in developing biomolecular therapies. Here, we show that encapsulation in protein cages formed by engineered variants of a cage-forming lumazine synthase establishes an effective route for microbial production and isolation of otherwise difficult-to-express, degradation-prone polypeptides. In this system, genetic fusion to a cage component protomer ensures efficient guest packaging while being produced in host bacterial cells. Meanwhile, the controlled opening outside the cellular context allows facile isolation of cargo via sequence-specific protease cleavage. Furthermore, modular patchwork assembly avoids guest overloading, preventing unwanted incomplete cage assembly and the formation of insoluble aggregates. The general applicability of our “encapsidic” production approach was demonstrated by the efficient production of six intrinsically disordered polypeptides that have proven therapeutic potentials.

## Introduction

Encapsulation in hollow proteinaceous compartments is a common biological strategy for the protective storage and delivery of otherwise unstable substances. Spatial segregation by physical barrier circumvents unwanted chemical transformation of cargoes, and their specialized release mechanism liberates them at the desired timing and location. For instance, viruses use capsids for protectively packaging their genomic materials and delivering them to the next host cells^1, 2, 3^. Ferritin stores iron ions in the lumen while preventing the production of toxic reactive oxygen species and releases them when needed^4^. These naturally occurring protein cages not only provide inspiration but also “almost-ready-to-use” platforms for developing customized nanometric molecular carriers^5, 6^.

Engineered protein cages have been exploited for both intra- and extracellular delivery routes. The majority of the current research seeks to load protein cages with molecules of interest outside the biological context to deliver the guest cargo into cells^7^. In the opposite scenario, protein cages can capture target molecules while being produced in host cells and transport them outside of the cells. For instance, viral capsids and their mimicries have been shown to package target RNA within cells and protect them from nucleases and other processing enzymes^8, 9, 10, 11, 12^. Such a protective storage function highlights the potential of protein cages as powerful tools for the intracellular production of valuable molecules that are otherwise rapidly degraded or converted by intracellular machinery.

Protein cages can protect encapsulated cargo proteins from degradation by cellular proteases. This has been demonstrated with cargo proteins carrying a degradation sequence in combination with a localization signal^13, 14, 15, 16, 17, 18^. While the degron tag would trigger the rapid proteolysis, encapsulation in protein shells mediated by the localization peptide prevents the degradation process by shielding cargoes from the proteolytic machinery. We have described such a rescue effect for engineered variants of a cage-forming lumazine synthase from *Aquifex aeolicus* (AaLS), where the negatively charged lumen of the shell-like structure protectively encapsulates a positively supercharged green fluorescent protein, GFP(+36), fused to a bacterial degradation tag^19^. However, in these previous studies, degradation signals have been used to demonstrate cargo encapsulation in the intracellular environment, and major questions and concerns have remained about the practicability of protein cages for heterologous protein production and isolation. Here, we report that an AaLS-based cargo encapsulation system offers a simple, robust, and efficient solution for the production of otherwise degradation-prone polypeptides in bacterial cells and their efficient isolation outside cellular contexts.

## Results

### Rescue of degradation-prone GFP by encapsulation in AaLS cages

We previously established a robust protein encapsulation system using a circularly permuted variant of AaLS (cpAaLS)^20^, which positions the N- and C-termini pointing towards the cage interior. In this system, a protein of interest is genetically fused to cpAaLS and simultaneously produced with an excess of another AaLS variant, such as wildtype AaLS (AaLS-wt), in *Escherichia coli* cells (Figure 1a). Coassembly of these AaLS proteins allows the formation of cage-like structures encapsulating the fusion partner in the lumen. The intracellular production level of these constituent proteins can be regulated individually by tetracycline or isopropyl-β-D-thiogalactopyranosid (IPTG). This is crucial to avoid excessive cargo loading that inhibits cage assembly due to steric hindrance^20^. This simple and modular compartmentalization system represents an ideal platform for exploring the potential of protein cages to produce degradation-prone proteins and peptides within cells. We refer to this heterologous protein production in capsid-like protein shells as “encapsidic” production.

**Figure 1.**
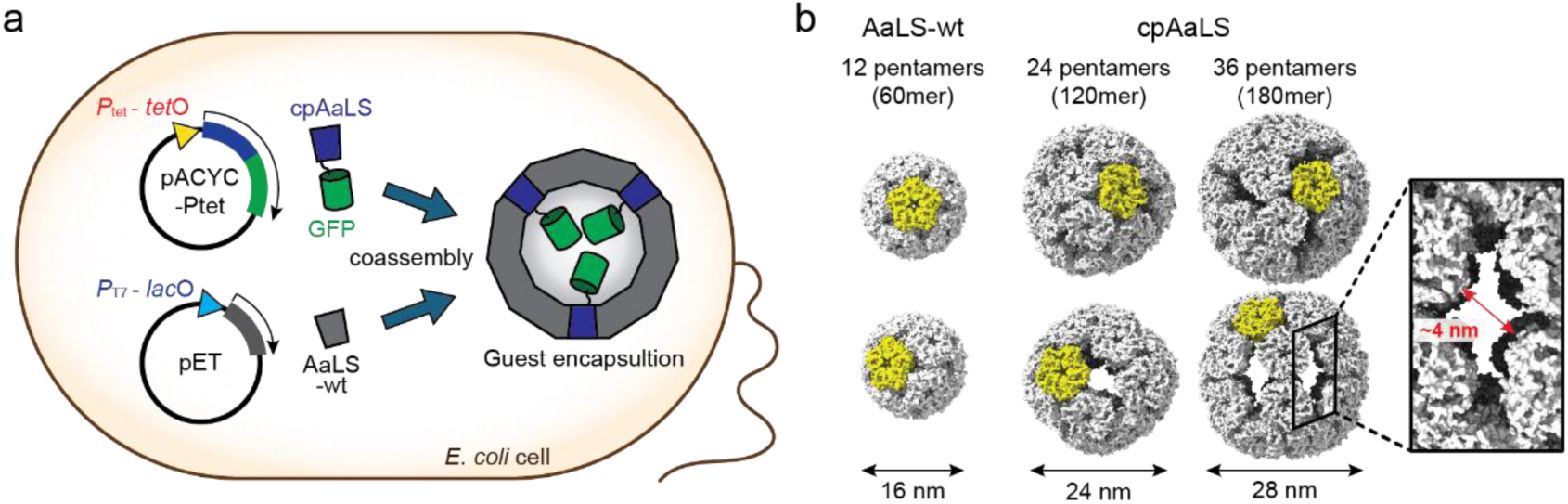
Guest encapsulation in AaLS cages using genetic fusion and patchwork assembly. (a) Schematic representation of the guest encapsulation. Coproduction of guest protein, e.g., GFP, fused to cpAaLS with another AaLS protein, e.g., AaLS-wt, in *E. coli* cells, leads to simultaneous coassembly and guest encapsulation in patchwork cages. Expression of the genes encoding these proteins is regulated individually by the tetracycline promoter combined with the tetracycline operator (*P*_tet_ – *tet*O) or the T7 promoter combined with the lactose operator (*P*_T7_ – *lac*O), respectively. (b) Structures of AaLS-wt (PDB ID: 1HQK) and cpAaLS (9G3O and 9G3N) assemblies. These cages are shown from two different viewing angles, and a representative pentamer unit is highlighted in yellow. Both cpAaLS assemblies have large keyhole-shaped pores in the wall, shown in a cropped and enlarged image.

AaLS protein is known to adopt a variety of assembly statuses upon mutagenesis^10, 21, 22, 23^, and the variant that predominantly constitutes patchwork cages rules the entire morphology^8, 20^. While the wildtype protein forms ∼16-nm dodecahedral structures composed of 12 copies of pentameric subunits (Figure 1b, left)^24^, the negatively supercharged variants, AaLS-neg and AaLS-13, assemble into expanded ∼28-nm and ∼40-nm particles composed of 36 and 72 pentamers, respectively^21^. When cpAaLS is coproduced with excess AaLS-wt, AaLS-neg, or AaLS-13, the morphology of the resulting patchwork structures resembles those formed by the latter AaLS variants alone^20^. In this study, we used AaLS-wt or a cpAaLS derivative possessing a His-tag in the linker connecting native termini (cpAaLS(His))^20^ as a scaffold protein to form patchwork cages. This cpAaLS(His) variant assembles into ∼24-nm and ∼28-nm hollow particles^20^ composed of 24- or 36-pentameric subunits, respectively. Notably, these expanded cage-like structures have large ∼4 nm keyhole-shaped pores in the walls (Figure 1b, middle and right)^22^. These AaLS variants with distinct assembly characteristics allow us to investigate the impact of the morphological features on the protection of encapsulated cargoes.

As a model of a degradation-prone cargo protein, we primarily employed a destabilized variant of GFP^19^. This variant possesses a bacterial signal peptide, SsrA^25^, which triggers the degradation of the fusion partner by the housekeeping proteases ClpXP and ClpAP^26^. When GFP fused to the SsrA tag (GFP-SsrA) is produced in *Escherichia coli* cells, the intracellular concentration of the protein remains low due to its rapid proteolysis. This expected effect was confirmed by flow cytometry analysis of *E. coli* producing GFP-SsrA and GFP without the tag (Supplementary Figure 1).

Encapsulation in protein cages should avoid a destined degradation of SsrA-tagged proteins by preventing their access to the proteolytic machinery^19^. To test this, GFP-SsrA was fused to the C-terminus of cpAaLS (cpAaLS-GFP-SsrA) and produced in *E. coli*. Notably, this fusion variant itself is unable to form an intact cage-like structure due to the steric hindrance of cargo GFP (Figure 2a)^20^. Indeed, flow cytometry analysis of cells producing cpAaLS-GFP-SsrA showed that the fluorescent intensity is comparable to background autofluorescence (Figure 2b,c, dark red and grey), indicating rapid degradation. However, coproduction of cpAaLS-GFP-SsrA with AaLS-wt or cpAaLS(His) increased the mean fluorescent intensity 73 or 63 times, respectively (Figure 2b,c, blue and orange), suggesting protective guest encapsulation for both combinations. Of note, these fluorescent values correspond to approximately 60% or 50% of the respective signals observed with cpAaLS-GFP lacking the SsrA tag (Figure 2b,c, red, orange, and blue). These rescue effects are significantly higher in comparison to a complimentary charge-driven encapsulation system we previously investigated^19^. Compared to reversible, non-covalent interaction, genetic fusion to a cage component protein probably yields a kinetic advantage for reducing cargo release and exposure to cellular proteases, resulting in increased levels of cargo protection.

**Figure 2.**
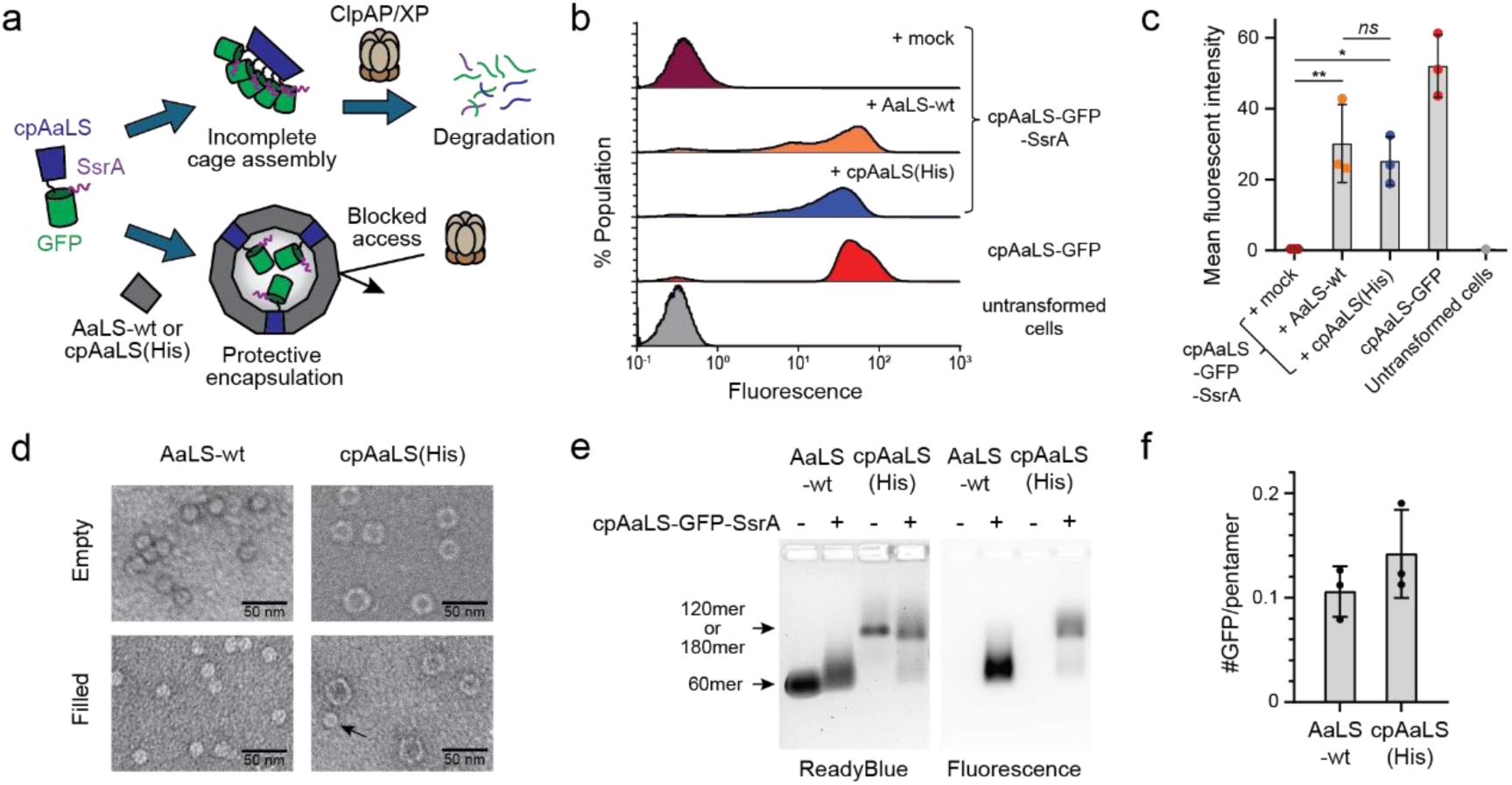
Protective encapsulation of a degradation-prone GFP variant. (a) Conceptual scheme of protective encapsulation of GFP equipped with a bacterial degradation tag SsrA (GFP-SsrA). When cpAaLS-GFP-SsrA is produced alone, cage assembly is hindered by guest steric repulsion, resulting in exposure of the SsrA tag and degradation by ClpAP/XP proteases. Patchwork coassembly with AaLS-wt or cpAaLS(His) proteins allows intact cage formation and guest protection. (b) Histogram representing fluorescence intensity within populations of cells coproducing cpAaLS-GFP-SsrA with either AaLS-wt (orange) or cpAaLS(His) (blue). The results from cells producing cpAaLS-GFP-SsrA with the mock plasmid (dark red), cpAaLS-GFP (red), and untransformed cells are also provided for comparison. 100% population corresponds to the total number of the analyzed cells (Y-axis scale, 0-1.2%). (c) Mean fluorescent intensity of the flow cytometry experiments. (d, e) Negative-stain transmission electroscope (TEM) images (c) and native-agarose gel electrophoresis (d) of AaLS-wt and cpAaLS(His) coproduced with cpAaLS-GFP-SsrA. The arrow in the microgram indicates a wildtype-like assembly of cpAaLS(His). The bands in the gel are visualized with ReadyBlue staining (left) and fluorescent (right). Bands corresponding to these AaLS cages are indicated. (f) Quantification of the number of GFP per pentamer. For panels c and f, data are shown as means ± standard deviation from three biological replicates. *ns* – *p* > 0.05; * – *p* < 0.1; ** – *p* < 0.01 (one-way ANOVA with Dunnett’s test).

The AaLS-wt and cpAaLS(His) proteins coproduced with cpAaLS-GFP-SsrA were isolated using affinity chromatography and characterized outside cells (Figure 2c-e). Negative-stain transmission electron microscopy (TEM) and native agarose gel electrophoresis (AGE) of the AaLS-wt sample confirmed successful complex formation with cpAaLS-GFP-SsrA resembling the observed morphology of the cages without cargo (Figure 2d,e). Efficient complex formation was also observed with cpAaLS(His) coproduced with cpAaLS-GFP-SsrA, where most of the particles were confirmed to be ∼24- or ∼28-nm spheres, but a small fraction of the wildtype-like ∼16-nm particles, that were absent with cpAaLS(His) alone, could also be detected (Figure 2d,e). Encapsulated GFP-SsrA and associated impurities (Supplementary Figure 2) possibly have an effect on the cpAaLS(His) assemblies. The number of GFP per AaLS-wt 60mer or cpAaLS(His) 180mer cage was estimated by the absorbance ratio at 280/475 nm as 1.3 and 5.1, corresponding to 0.11 and 0.14 GFP molecules per pentamer, respectively (Figure 2f)^22, 27^. Together with the presented flow cytometry experiments, these results confirm little difference between the AaLS-wt and cpAaLS(His) cages in protective guest encapsulation of the SsrA-tagged proteins.

### Cargo GFP retrieval by protease cleavage

Upon establishing a system that successfully protects the degradation-prone GFP variant from cellular proteases, we next sought to retrieve the cargo proteins from the AaLS cages and redesigned the protein constructs accordingly. First, a recognition sequence for the commonly used tobacco etch virus TEV protease^28, 29, 30^ was introduced between cpAaLS and GFP-SsrA, yielding cpAaLS-tev*-GFP-SsrA (Figure 3a), so that the cargo proteins can be disconnected from the cage-forming protomers by proteolytic cleavage. Additionally, the His-tag region of AaLS(His) was replaced by a Strep II tag as nickel-based affinity chromatography is unsuitable for purifying therapeutic biomolecules due to the potential contamination of toxic metal ions (Supplementary Table 1). This variant, called cpAaLS(Strep), was produced as a mixture of 24- and 28-nm spherical assemblies, like the parent cpAaLS(His) (Supplementary Figure 3). Notably, this tag replacement and the change in affinity chromatography method resulted in a significantly improved purification yield: ∼1 mg cpAaLS(His) and ∼15 mg cpAaLS(Strep) from *E. coli* cell culture in a 0.25-L Luria-Bertani (LB) medium.

**Figure 3.**
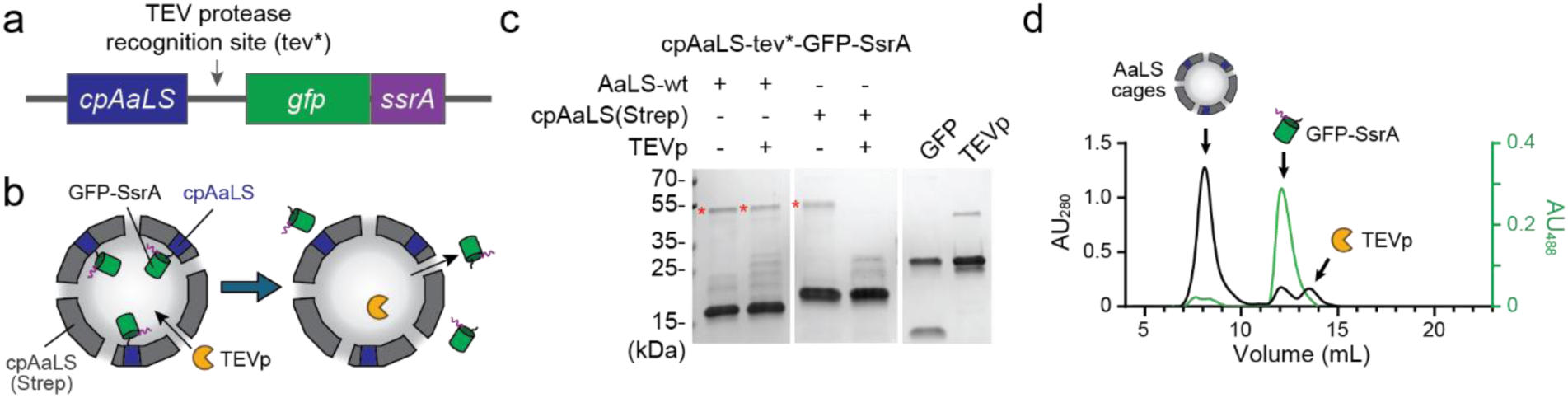
GFP retrieval from AaLS cages. (a) Design of cpAaLS fusion to GFP-SsrA via TEV protease recognition site (tev*). (b) Hypothetical mechanisms of TEV protease (TEVp) entry into cpAaLS(Strep) cages through the ∼4-nm pores and resulting GFP release. (c) SDS-PAGE analysis of the cleavage reaction. Red asterisks indicate the bands corresponding to cpAaLS-tev*-GFP-ssrA (47.6 kDa). The theoretical molecular masses of other proteins are as follows: AaLS-wt, 18.1 kDa; cpAaLS(Step), 18.5 kDa; TEVp, 28.8 kDa; and GFP-SsrA, 29.1 kDa. The bands for TEVp and GFP-SsrA are undistinguishable due to the minor differences in their molecular masses. (d) Size-exclusion chromatography confirming that GFP-SsrA is released from the cpAaLS cages upon cleavage, shown as absorbance at 280 nm (black) and 488 nm (green). AaLS cages were eluted at the void volume (7-8 mL) of a Superdex 75 column.

TEV protease (∼3.5 nm) is smaller than the pores (∼4 nm) of cpAaLS assemblies (Figure 1b). Therefore, we assumed the enzyme would be able to enter the inner cavity of the protein cages and release the cargo GFP by proteolytic digest (Figure 3b), while this wouldn’t occur with the AaLS-wt cages possessing only small (∼3 Å) pores (Figure 1b). To verify this hypothesis, AaLS-wt and cpAaLS(Strep) patchwork assemblies containing cpAaLS-tev*-GFP-SsrA were treated with an engineered variant of TEV proteases^28, 29, 30^, and the cleavage was monitored by SDS-PAGE (Figure 3c). Unexpectedly, the proteolytic reaction was observed for the cages based on AaLS-wt, albeit at a low efficiency of ∼30%. This partial cleavage might result from imperfectly assembled AaLS-wt cages, possibly due to overloading by cargo. Strikingly, the cleavage efficiency for patchwork cages based on cpAaLS(Strep) was almost complete. These results underline that cage-like assemblies with large pores enable the TEV protease to enter the protein cages and cleave the cargo at the specific recognition motif (Figure 3b). This cleavage does not lead to a noticeable morphology change for either type of AaLS cages, as judged by the TEM images (Supplementary Figure 4). Size-exclusion chromatography (SEC) confirmed that the cleaved GFP was indeed released from the still intact host cages (Figure 3d).

The keyhole-shaped, ∼4-nm dimension pores of cpAaLS(Strep) cages likely allow passage of smaller molecules (Figure 1 and Supplementary Figure 5a). This is advantageous for guest retrieval using a sequence-specific protease such as TEV protease and guest release upon cleavage in vitro. At the same time, this porous shell still serves as a barrier to prevent the internalization of major proteolytic machinery in bacterial cells such as ClpXP or FtsH that are much larger (∼16-nm) or are membrane-bound (Supplementary Figure 5b)^31^. Therefore, the porous nature of cpAaLS(Strep) cages appears to provide an ideal system for microbial production of degradation-prone proteins and their retrieval outside cellular contexts.

### Encapsidic production of a glucagon-like peptide 1 derivative, lixisenatide

Beyond the SsrA-based proof-of-concept model, we envisaged that the encapsidic production system would be useful for clinically relevant bioactive molecules. Along this line, we focused on short, intrinsically disordered peptides that are typically difficult to produce recombinantly due to the rapid degradation by endogenous proteases in host cells^32, 33^. The current peptide production largely relies on chemical synthesis, which requires excess molarity of expensive and harmful reactants and, therefore, remains a concern in costs and waste management^34^. Furthermore, synthesizing relatively long peptides, >40 residues, is often burdensome, whereas amino acid length is ultimately not a limiting factor in recombinant protein production. Given the remarkable progress in peptide therapeutic development in the last decade and its increasing global market^35, 36^, there is a need for an inexpensive and ecological platform for their production, where the microbe-based encapsidic production system potentially meets the criteria.

As an initial model of therapeutic peptides, we selected a glucagon-like peptide 1 (GLP-1) derivative, lixisenatide (LIX), an antidiabetic drug composed of 44 amino acids (Supplementary Table 2)^37^. This peptide was fused to the C-terminus of cpAaLS via a TEV protease recognition site, referred to as cpAaLS-tev*-LIX, and coproduced with cpAaLS(His) or cpAaLS(Strep) (Figure 4a, Patchwork cage). Additionally, we tested a sole production of cpAaLS(His) or cpAaLS(Strep) fusion to LIX, cpAaLS(His)-tev*-LIX or cpAaLS(Strep)-tev*-LIX, respectively (Figure 4a, Single-component cage), since such a small peptide might not hinder cage assembly. For comparison, the peptide was also produced in fusion with His-tagged maltose-binding protein (MBP-LIX), frequently used as a fusion partner for microbial production of short peptides^38^, as well as with His-tag only (LIX). Those constructs were produced in *E. coli* cells and analyzed by SDS-PAGE using whole cells and soluble fractions after bacteriolysis (Figure 4b,c and Supplementary Figure 6). While the His-tagged LIX was undetectable, substantial bands corresponding to the LIX fusion forms were observed for all other cases. The highest yield in the soluble fraction was obtained with the sole production of cpAaLS(His)-tev*-LIX, followed by its Strep-tagged variant cpAaLS(Strep)-tev*-LIX. These results suggested that the patchwork assembly approach is only necessary for bulky proteins and is not required for the small LIX peptide production within the AaLS cages. Furthermore, the encapsidic production seems superior to the conventional MBP-fusion strategy in terms of total production level and solubility. Considering the Strep-tagged cpAaLS showed a 15-fold higher purification yield than the His-tagged variant, we conclude that cpAaLS(Strep)-fusion is the best form among other tested constructs for producing LIX in *E. coli*.

**Figure 4.**
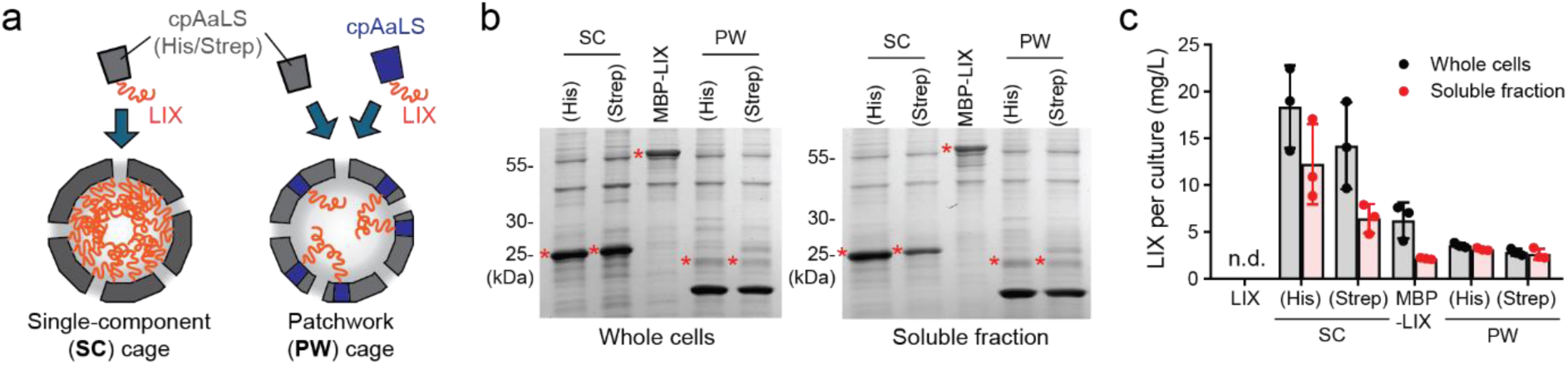
Encapsidic production of lixisenatide. (a) Cartoon illustration of single component (SC) and patchwork (PW) cages containing lixisenatide (LIX). (b, c) SDS-PAGE analysis for LIX production using whole cells (left gel) and soluble fraction after bacteriolysis (right gel). The labels (His) or (Strep) indicate the purification tag of cpAaLS variants used for the single component (SC) or patchwork (PW) cages. LIX, LIX peptide alone; MBP-LIX, LIX fused to maltose-binding protein. The bands indicated with red asterisks were used for densitometry assay to estimate the LIX yield per litter of cell culture (c). Calculated molecular masses of proteins are; LIX, 6.6 kDa; cpAaLS(His)-LIX, 24.3 kDa; cpAaLS(Strep)-LIX, 24.6 kDa; MBP-LIX, 47.3 kDa; cpAaLS-LIX, 23.6 kDa. cpAaLS(His), 18.2 kDa; and cpAaLS(Strep), 18.5 kDa. Uncropped gel images, including BSA quantification standards, are also provided in Supplementary Figure 6. For panel c, data are shown as means ± standard deviation from three biological replicates. n.d., not detected.

### Encapsidic production of other peptides with therapeutic potential

The general applicability of the encapsidic production system was investigated using five other peptides possessing therapeutic potential: Teriparatide (TER)^39^, Aviptadil (AVI)^40^, Thymosin β-4 (THY)^41^, Enfuvirtide (ENF)^42^, and Magainine-2 (MAG)^43^ (Supplementary Table 2). They were analogously produced as a fusion to cpAaLS(Strep) using *E. coli* cells and analyzed by SDS-PAGE. However, this showed that only THY was produced in a soluble form, and others almost exclusively ended up in insoluble fractions upon bacteriolysis (Figure 5a,b and Supplementary Figure 7). In the case of MAG, substantial cytotoxicity was also observed, likely due to the bacteriolytic activity. These peptides that showed poor solubility with the encapsidic system are more hydrophobic than LIX and THY (Supplementary Table 2), likely facilitating their aggregation and insolubility.

**Figure 5.**
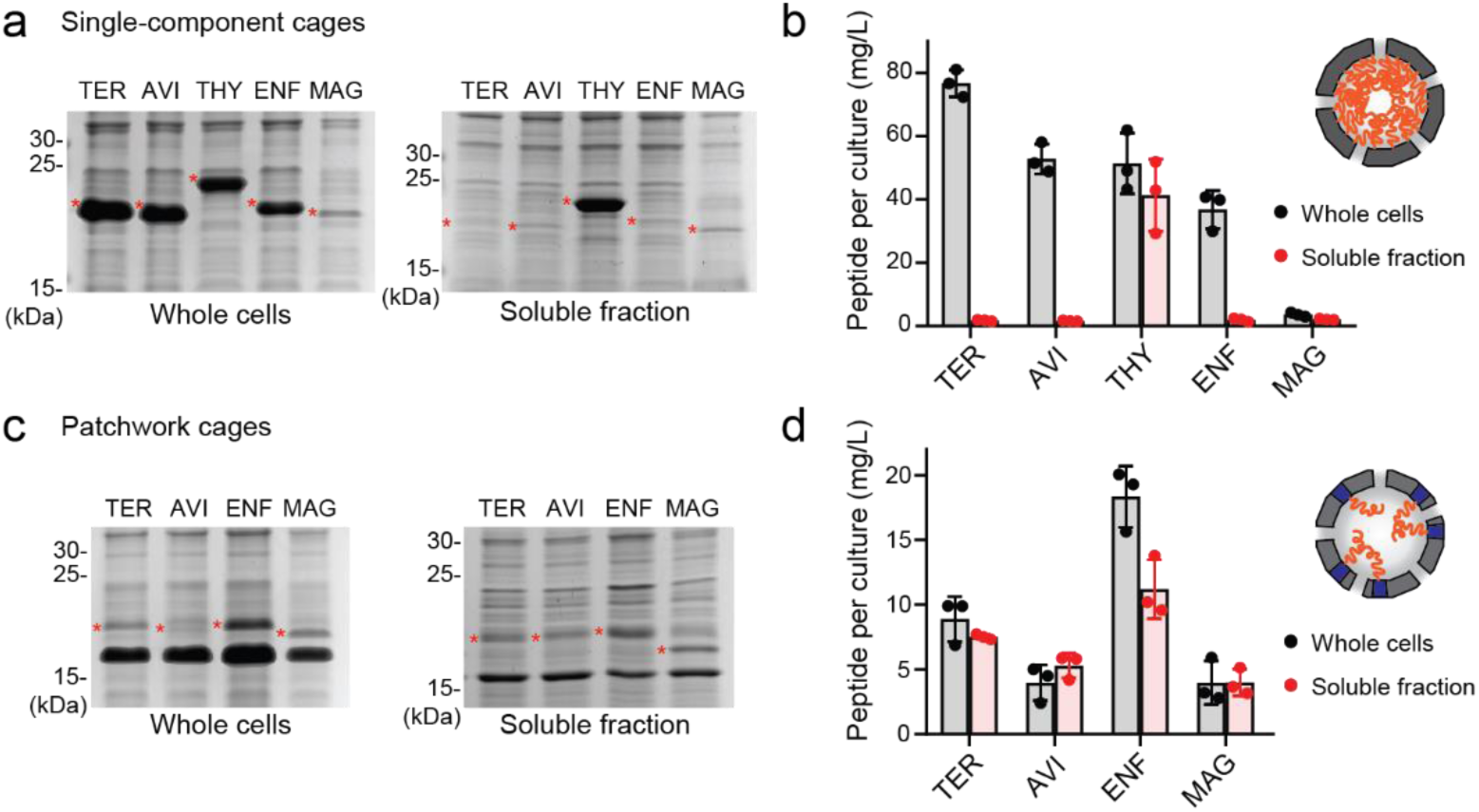
Encapsidic production of therapeutic peptides. (a-d) SDS-PAGE analysis for peptides produced in single-component cages (a,b) or patchwork cages (c,d) using whole cells (a,c, left gel) and soluble fraction after bacteriolysis (a,c, right gel). The bands indicated with red asterisks were used for densitometry assay to estimate the yield per litter of cell culture (b,d). Calculated molecular masses of proteins are; cpAaLS(Strep)-TER, 24.0 kDa; cpAaLS(Strep)-AVI, 23.1 kDa; cpAaLS(Strep)-THY, 19.8 kDa; cpAaLS(Strep)-ENF, 24.2 kDa; cpAaLS(Strep)-MAG, 22.2 kDa; cpAaLS-TER, 22.8 kDa; cpAaLS-AVI, 22.0 kDa; cpAaLS-ENF, 23.1 kDa; cpAaLS-MAG, 21.2 kDa. Uncropped gel images, including BSA quantification standard, are also provided in Supplementary Figures 7 and 8. For panels b and d, data are shown as means ± standard deviation from three biological replicates.

Reducing the fusion content of the protein cage using the patchwork assembly can overcome the solubility issues of the hydrophobic guests. These peptides were fused to cpAaLS and coproduced with cpAaLS(Strep). At 0.4 µg/mL tetracycline, nearly all the produced fusion protein was obtained in a soluble form for TER, AVI, and MAG, and ∼60% for ENF (Figure 5c,d). Unlike the single-component cage strategy with MAG, no remarkable cytotoxicity was observed for any peptides. Induction with higher tetracycline concentration resulted in an increased proportion of the insoluble fraction for all the cases and no improvement in the protein yield in the soluble fraction (Supplementary Figure 8). Therefore, we conclude that optimization of guest density is crucial for producing hydrophobic peptides within protein cages in a soluble form. Furthermore, these results highlight that the patchwork assembly strategy is advantageous not only for intact cage formation with bulky protein cargo but also for solubilizing aggregation-prone peptides.

### Isolation and characterization of encapsidically produced lixisenatide

After the successful encapsidic production of the LIX peptide, we tested the guest retrieval from the protein cages. Similarly to the GFP system, isolated cpAaLS(Strep)-tev*-LIX protein was treated with TEV protease. However, the cleavage was barely observed under a buffer condition used for GFP retrieval, 50 mM Tris-HCl buffer (pH 8.0) containing 0.3 M NaCl (Figure 6a). Even though the cpAaLS(Strep) cages have pores, the high density of the guest peptide likely limits access of TEV protease to the lumen.

**Figure 6.**
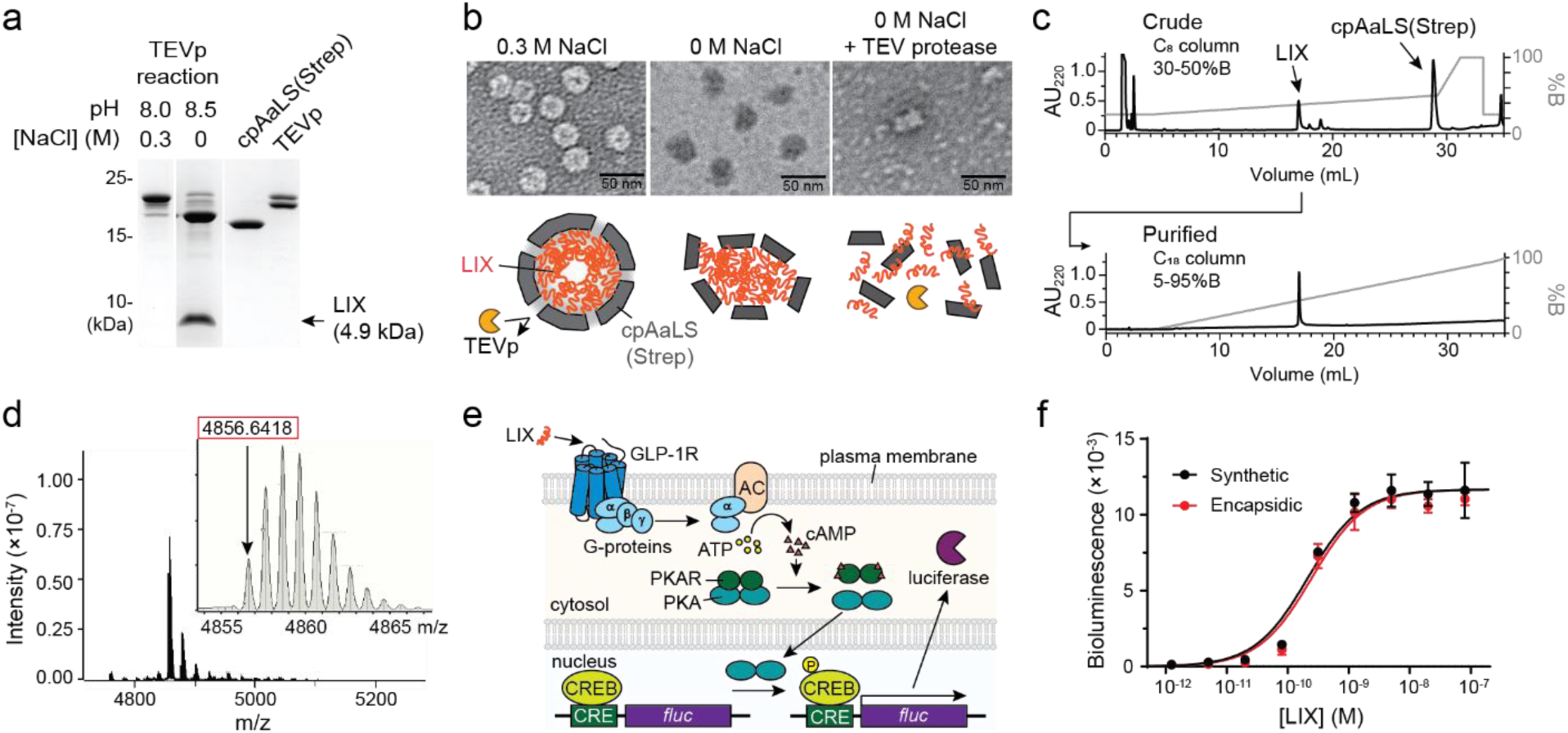
LIX retrieval and characterization. (a) SDS-PAGE analysis of cpAaLS(Strep)-tev-LIX treated by TEV protease in 50 mM Tris-HCl buffers (pH 8.0 or 8.5) containing 0.3 or 0 M NaCl, respectively. The arrow indicates the band corresponding LIX (4.9 kDa). The theoretical molecular masses of other proteins are as follows: cpAaLS(Strep)-tev-LIX, 24.6 kDa; cpAaLS(Step), 18.5 kDa; TEVp, 28.8 kDa. The corresponding uncropped gel image is provided in Supplementary Figure 9. (b) Negative-stain TEM images of cpAaLS(Strep)-tev-LIX in 50 mM Tris-HCl buffer (pH 8.0) containing 0.3 M NaCl (left) and in 50 mM Tris-HCl buffer (pH 8.5) before (middle) and after (right) TEV protease treatment. A possible mechanism of TEV protease access to the guest peptide is shown below the micrographs. (c) Reverse-phase high-performance liquid chromatography (RP-HPLC) analysis of crude (top) and purified LIX (bottom). (d) Mass spectrometry of purified LIX. The enlarged view of the main peak is shown in the inset. The theoretical monoisotopic mass of LIX is 4856.53. (e) Diagram of the glucagon-like peptide-1 receptor (GLP-1R)-mediated reporter assay system. AC, adenylnyl cyclase; PKA, protein kinase A; PKAR, PKA regulator; CRE, cAMP-response element; CREB, CRE-binding protein; fluc, firefly luciferase; P, phosphate group. (e) The activity of LIX on stimulation of GLP-1R. Data using a commercially available synthetic standard (black) are provided as a reference for the encapsidically produced and isolated LIX (red). Data are shown as means ± standard deviation from triplicate measurements. EC_50_ of synthetic and encapsidically produced LIX were determined to be 0.28 ± 0.09 and 0.35 ± 0.15 nM, respectively.

Efficient protease cleavage to release the LIX peptide from cpAaLS(Strep) was achieved by changing the reaction buffer to 50 mM Tris-HCl buffer (pH 8.5) (Figure 5a). This low ionic strength, alkaline condition was previously found to disassemble cpAaLS cages into the pentameric subunits^22^, which was also the case for the cpAaLS(Strep) variant (Supplementary Figure 2b). While the TEM images of the isolated cpAaLS(Strep)-tev-LIX in 50 mM Tris-HCl (pH 8.0) buffer containing 0.3 M NaCl showed regular near-spherical structures with approximately 24-nm size (Figure 5b, left), the same protein in 50 mM Tris-HCl buffer at pH 8.5 exhibited a similar size of particles but in uneven shapes (Figure 5b, middle), likely reflecting partially broken shells. After the TEV protease cleavage reaction, they mostly disassembled into smaller fragments, and cage-like structures were not observed (Figure 5b, right). The efficient cleavage was also observed in 50 mM Tris-HCl buffer at pH 7.5 and 8.0 without additional salt, but did not occur with 0.1-0.3 M NaCl regardless of pH (Supplementary Figure 9). These results suggest that destabilization of cpAaLS(Strep) assembly by lowering ionic strength is crucial for retrieval of the cargo peptides by TEV protease cleavage.

After the cleavage reaction, the sample containing LIX peptide was passed through Strep-Tactin affinity chromatography to remove cpAaLS(Strep) and the TEV protease, which also possesses a strep-tag. Subsequent purification using reverse-phase high-performance liquid chromatography (RP-HPLC) yielded the LIX peptide with ∼100% purity (Figure 5c). The fidelity of the purified LIX was confirmed by electrospray ionization mass spectrometry (ESI-MS), observing 4856.64 (theoretical mass: 4856.53) (Figure 6d and Supplementary Figure 10). Approximately 0.3 mg of LIX was obtained from an *E. coli* cell culture in a 0.5-L LB medium.

The activity of encapsidically produced LIX peptide was tested using a genetically engineered human embryonic kidney (HEK) cell line that produces human GLP-1 receptor (GLP-1R) (Supplementary Figure 11) and carries a firefly luciferase reporter under the control of cyclic adenosine monophosphate (cAMP) response element (CRE) (Figure 6e). The binding of LIX to the GLP-1R should trigger the activation of coupled G-proteins, leading to an increase in intracellular concentrations of cAMP. Bioluminescence of luciferase, of which the gene expression is induced by cAMP, was used as a readout of the successful stimulation of the GLP-1R. The encapsidically produced LIX peptide showed GLP-1R stimulation activity comparable to a synthetic standard of the same peptide (Figure 6f). These results exemplify that the encapsidic production can be utilized for a functional therapeutic peptide.

## Discussion

The subcellular compartmentalization using an engineered protein cage establishes a practical means for microbial production and isolation of degradation-prone polypeptides. Genetic fusion to a cage-component protein ensures efficient compartmentalization of cargo peptides, protecting them from cellular proteolytic machinery. Besides proteolysis focused here, spatial segregation by protein shells potentially circumvents other problems encountered during heterologous protein production. For instance, cytotoxicity of a membrane-disrupting antimicrobial peptide MAG can be suppressed by encapsulation in AaLS protein cages (Figure 5)^43^, while a similar attenuation effect has also been shown for a toxic protease from human immunodeficiency virus (HIV)^44^. Protein shells may also prevent the entry of undesired cellular enzymes, such as reductases that cleave disulfide bonds and destabilize protein folding^45^. The patchwork assembly system is advantageous for these purposes, as it allows customizing the morphology and chemical properties of protein cages for individual cargo by selecting an appropriate variant as a scaffolding protein^20^. For example, protein cages formed by negatively supercharged AaLS variants can be utilized to exclude unwanted polyanionic molecules^46^. This modularity makes the AaLS-based system a potent technology for producing a variety of difficult-to-express proteins.

To date, substantial efforts have been directed towards the efficient microbial production of short peptides as an ecological and economical alternative to canonical chemical synthesis. As shown with MBP for LIX (Figure 4b), fusion to a soluble protein can reduce the degradation and cytotoxicity, the most common issues during recombinant peptide production^38^. However, this approach still exposes peptides, leaving a chance for proteolysis during production and purification. An alternative strategy is based on the targeting of peptides to inclusion bodies that also segregate the products from other cellular contents^47^. Meanwhile, retrieval of aggregation-prone peptides from the inclusion body often requires harsh chemical conditions, including high concentrations of chaotropic agents, which limit the usage of enzymes for downstream processes, e.g., sequence-specific cleavage. Protein cages that solubilize and segregate cargo peptides until exposing them by lowering ionic strength provide an ideal scenario, enabling both protection within cells and further enzymatic processing of peptides.

Therapeutic peptide production within protein cages is potentially useful for drug discovery and delivery. This genetically encoded peptide biosynthesis can facilitate their mutagenesis and evolutionary engineering toward improved biological activity, e.g., higher affinity to target receptors. As shown for lixisenatide, affinity chromatography followed by protease cleavage yielded a desired peptide with purity likely sufficient for screening purposes (Figure 5c). Furthermore, protein cages that apparently enlarge and shield cargo molecules may improve otherwise limited bioavailability of peptide therapeutics that are, in general, rapidly eliminated by renal filtration and degraded by proteases because of their small and flexible structural characteristics^35^. On the other hand, the cargo release, which relies on lowering ionic strength and a virus-derived protease, is most likely unsuitable for direct in vivo application. Toward the development of more versatile, smart nanocarriers, further reengineering of AaLS protein to induce cage disassembly and cargo release by a defined stimulus of interest is ongoing in our laboratory.

## Materials and Methods

### Plasmids

Plasmid pET15 derivatives were used for production of AaLS-wt, cpAaLS(His), cpAaLS(Strep), and cpAaLS(Strep) fused to model peptides, and pACYC-Ptet^48^ for cpAaLS fusion to GFP and peptides in the patchwork strategy. The detailed procedures for preparing these plasmids are provided in the Supplementary Information, where the protein, the plasmid, and the oligonucleotide sequences used in this study are summarized in Supplementary Tables 1, 3, and 4.

### Protein production

AaLS proteins were produced in *E. coli* strain BL21-gold (DE3) that were transformed with corresponding plasmids. After culturing the cells to OD_600_ of ∼0.7, protein production was induced by adding 0.2 mM isopropyl β-D-1-thiogalactopyranoside (IPTG), followed by further culturing cells at 25°C for ∼18 hours. For patchwork assembly, cpAaLS fusion production was induced with 0.4 μg/mL tetracycline unless specified.

### Flow cytometry

*E. coli* cells producing GFP were analyzed by flow cytometry, as previously described^19^. Briefly, cell cultures were diluted ∼300 fold with phosphate-buffered saline and subjected to measurements on a Navious Flow Cytometer (Beckman Coulter, Brea, CA, USA). A blue laser operating at 488 nm was used for excitation, and fluorescence was detected through a 525/40 nm emission filter. Data were analyzed using Kaluza C software (Beckman Coulter). Experiments were performed in triplicate, and one representative histogram (Figure 2a) and the mean fluorescent intensity of each data point, in addition to means ± standard (Figure 2b), is provided.

### Protein purification

AaLS protein, which possesses either a His6 or a StrepII tag, was isolated by affinity chromatography using HisPur Ni-NTA resin (Thermo Fisher Scientific, Waltham, MA, USA) or Strep-Tactin XT 4Flow resin (Iba Lifescience, Goettingen, Germany), respectively. After elution, the buffers were replaced with 50 mM Tris-HCl buffer (pH 8.0) containing 0.3 M NaCl and 5 mM ethylenediaminetetraacetatic acid (EDTA) using Amicon Ultra centrifugal units (50 kDa MWCO, Merck-Millipore, Burlington, MA, USA). The resulting solution was used as a stock for further analysis by SDS-PAGE, native agarose electrophoresis (AGE), absorbance spectrometry, and transmission electron microscopy (TEM). AaLS concentration was determined by absorbance at 280 nm using the extension coefficients listed in Supplementary Table 1.

### Native agarose gel electrophoresis

Samples were prepared in phosphate-buffered saline containing 1 mM EDTA (PBS-E) and added with a 4× native-PAGE loading buffer (0.2 M BisTris-HCl buffer (pH 7.2) containing 40% (v/v) glycerol, 0.016% (w/v) bromophenol blue). Electrophoresis was performed using 1.0% (w/v) agarose gels and a running buffer (0.2 M Tris base, 25 mM tricine, pH adjusted with phosphoric acid to ∼7.6) at 4 °C and 70 V for 90 min. Fluorescent bands were visualized using a ChemiDoc MP (Bio-Rad, Hercules, CA, USA) in the fluorescein detection mode (460–490 nm excitation and 532/28 nm emission filters). The same gels were stained with ReadyBlue Protein Gel Stain (Sigma-Aldrich, St. Louis, MO, USA).

### Negative-stain transmission electron microscopy

Approximately 0.035 mg/mL of protein solution was placed on a glow-discharge-treated carbon-coated copper grid (EM resolution, Sheffield, UK). Excess solution was removed using filter paper followed by staining with UranyLess EM Stain (Electron Microscope Sciences, Hatfield, PA, USA). Samples were visualized on a JEM-1230 electron microscope with 80 kV operation (JEOL ltd., Tokyo, Japan). Obtained images were analyzed using Image J software (National Institutes of Health, Bethesda, MD, USA).

### SDS-PAGE

Samples were prepared in an SDS-PAGE sample buffer (62.5 mm Tris-HCl buffer (pH 6.8) containing 2.5% (w/v) sodium dodecyl sulfate (SDS), 10% (v/v) glycerol, 5% (v/v) β-mercaptoethanol, 0.2 mM dithiothreitol (DTT), and 0.004% (w/v) bromophenol blue) and heated at 95 °C for 10 min. Electrophoresis was performed using 2 µg proteins, 15% polyacrylamide gels, unless specified, and a Tris-Tricine buffer system at 80 V for 10 min followed by 150 V for 60 min. As a protein standard, PageRuler™ Prestained Protein Ladder (Thermo Fisher Scientific) was used. Protein bands were visualized using ReadyBlue Protein Gel Stain.

### GFP encapsulation efficiency

The number of encapsulated GFP was estimated based on the absorbance ratio at 280/475 nm, as previously reported^27^. Absorbance spectra were measured at 25 °C on a UV-1900 spectrophotometer (Shimadzu, Kyoto, Japan) using a 1-cm-light-pass quartz cuvette and protein in PBS-E. The calculation details are described in Supplementary Information.

### GFP retrieval

The cpAaLS-tev-GFP-ssrA assemblies with AaLS-wt or cpAaLS(Strep) in 50 mM Tris-HCl buffer (pH 8.0) containing 0.3 M NaCl, 1 mM EDTA, and 2 mM DTT were added with super TEV protease^49^ possessing a Strep-tag (1:15 as mass ratio) and kept at room temperature overnight. The cleavage reaction and guest release were assessed by SDS-PAGE and size-exclusion chromatography using a Superdex increase 75 10/300 column (Cytiva Europe GmbH, Freiburg im Breisgau, Germany), respectively.

### Peptide production test

Protein production was performed using the above-described protocol in a 4-mL scale, and the samples corresponding to 8 µL of cell culture before and after sonication were loaded on SDS-PAGE gel. For densitometry analysis, bovine serum albumin (BSA, Sigma-Aldrich) was used as standard, and the band intensities were quantified on Image Lab software version 6.1 (Biorad). The peptide yield was estimated with the molecular mass ratio of peptide to the fusion protein and provided as the peptide yield per 1 L of cell culture using the following equation.

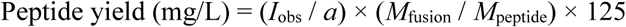

where *I*_obs_ is the observed band intensity, *a* is the correlation coefficient estimated from the BSA standard curve, and *M*_fusion_ and *M*_peptide_ are molecular masses of fusion protein and peptide, respectively. Experiments were performed in triplicate, and means ± standard deviation are provided.

### LIX retrieval

The cpAaLS(Strep)-LIX sample in 50 mM Tris-HCl (pH 8.5) containing 1 mM EDTA and 2 mM DTT was added with super TEV protease possessing a Strep-tag (1:15 as mass ratio) and kept at room temperature overnight. After reducing cpAaLS(Strep) and sTEVp contents by passing through StrepTactin XT resin, the crude peptide was subsequently purified by reverse-phase high-performance liquid chromatography (RP-HPLC) using a Nucleosil C8 column (120 Å, 5 μm, 4.6 × 150 mm) (Macherey Nagel, Düren, Germany). The purity and the fidelity of the peptide were confirmed by RP-HPLC using a Nucleosil C18 column (120 Å, 5 μm, 4.6 × 150 mm) and electron spray ionization time-of-flight mass spectrometry (ESI-TOF-MS) on a Bruker Daltonics micrOTOF-Q II (Billerica, MA, USA), respectively.

### Lixisenatide activity assay

The CRE/CREB luciferase reporter HEK293 cell line (BPS Bioscience, San Diego, CA, USA) was transduced with GLP1R (NM_002062) human tagged ORF clone lentiviral particle (OriGene, Rockville, MD, USA). Production of GLP1R was confirmed by flow cytometry using human GLP-1R Alexa Fluor 488-conjugated antibody (R&D Systems, Minneapolis, MN, USA) following the manufacturer’s protocol (Supplementary Figure 9). A detailed version of the cell culture, transduction, and antibody staining was provided in the Supplementary Information.

Luciferase reporter assay was performed using a Pierce firefly luciferase glow assay kit (Thermo Fisher Scientific) following the manufacturer’s protocol. Specifically, the genetically engineered CRE/CREB luciferase reporter HEK293 cells that produce GLP-1R were seeded in a 96-well microplate (2 × 10^4^ cells per well) and cultured in the Dulbecco’s modified Eagle’s medium with high glucose (DMEM-HG) supplemented with 10% heat-inactivated fetal bovine serum (FBS), 100 U/mL penicillin and 0.1 mg/mL streptomycin, referred to as complete media (CM), at 37°C overnight. The next day, the cells were treated with LIX samples (1.25 pM - 80 nM) in CM at 37°C for 16 h. After washing with Dulbecco’s modified phosphate-buffered saline (DPBS), cells were lysed with the Cell Lysis Buffer while gently shaking at 15°C for 30 min. Fifty microliters of the Firefly Glow Assay Buffer containing D-luciferin (0.6 mg/mL) in a black 96-well microplate was added with 20 µL cell lysate and incubated at room temperature for 10 min. Bioluminescence was monitored on an Infinite M200 PRO plate reader (Tecan, Männedorf, Switzerland). Experiments were triplicated each time, and the whole assay was repeated twice. The results from one repetition are shown in Figure 6. Half maximal effective concentration (EC_50_) was estimated by curve-fitting data points with the Hill equation using Graph Pad Prism software (La Jolla, CA, USA), and means ± standard deviations from two repeats are provided.

## Acknowledgments

This work is generously supported by the National Science Centre of Poland (OPUS-18 K/PBO/000754) and an EMBO Installation Grant. We would like to thank Dr. Michał Bochenek (Malopolska Centre of Biotechnology (MCB), Jagiellonian University (JU)) for his help with the flow cytometry experiments, Dr. Urszula Jankowska and Dr. Bożena Skupień-Rabian (MCB, JU) with mass spectrometry, as well as Dr. Olga Woźnicka and Dr. Elżbieta Pyza (Faculty of Biology, JU) with negative-stain electron microscopy. D.Kw is grateful for a European Research Area (ERA) fellowship.

## Author contributions

A.G., J.P., M.Z., D.Kw., D.Kl, and Y.A. performed experiments and analyzed data. Y.A. conceptualized the project, oversaw the research, acquired the funding, prepared the figures, and wrote the original draft. A.G., J.P., M.Z., D.Kw., L.K., S.G., N.KT., and Y. A. reviewed and edited the manuscript. All the authors contributed to the experimental design, as well as read and approved the final version of the manuscript.

## Additional information

Supplementary Information accompanies this paper at http://XXX

## Competing interests

The authors declare no competing financial interests.

## Notes

### Competing Interest Statement

The authors have declared no competing interest.

