## Supplementary Information for "Encapsidic production and isolation of degradation-prone polypeptides"

**This file includes:**

**Supplementary Figures 1-11**

**Supplementary Tables 1-4**

**Detailed Materials and Methods**

**Uncropped gel images 1-7**

#### Supplementary Figures

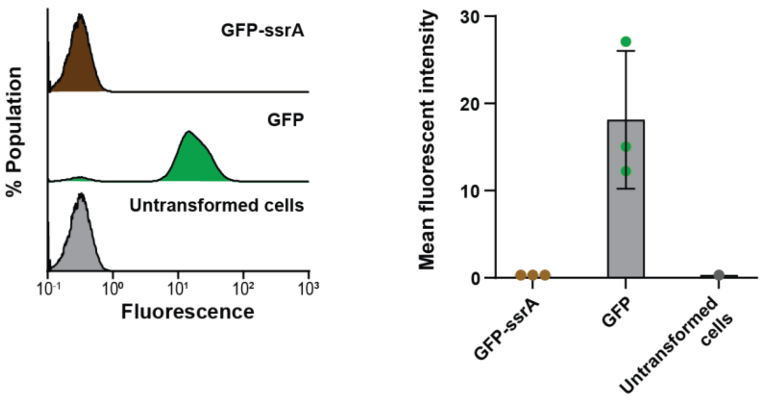


**Supplementary Figure 1. Flow cytometry analysis of cells producing GFP-SsrA.** Histogram (left) and mean (right) fluorescent intensity of *E. coli* cells producing GFP-SsrA (brown) or GFP (green). 100% population corresponds to the total number of the analyzed cells (Y-axis scale, 0-1.2%). The data for untransformed cells (grey) correspond to Figure 1b,c. The bar graph presents means ± standard deviation from three biological replicates.

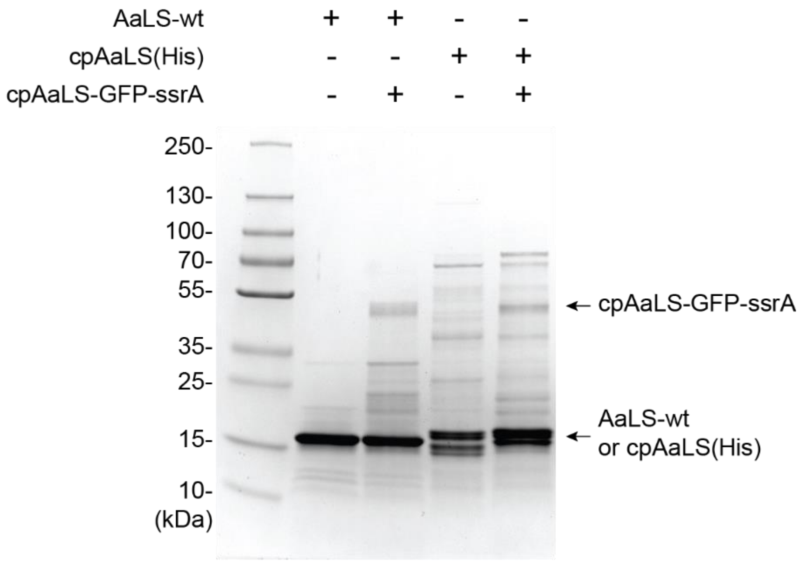


**Supplementary Figure 2. SDS-PAGE analysis of AaLS cages containing cpAaLS-GFP-ssrA.** Electrophoresis was performed using a 4-20% gradient acrylamide gel, followed by staining with ReadyBlue. Calculated molecular masses of proteins are; AaLS-wt, 18.1 kDa; cpAaLS(His), 18.4 kDa; and cpAaLS-GFP-ssrA, 46.7 kDa.

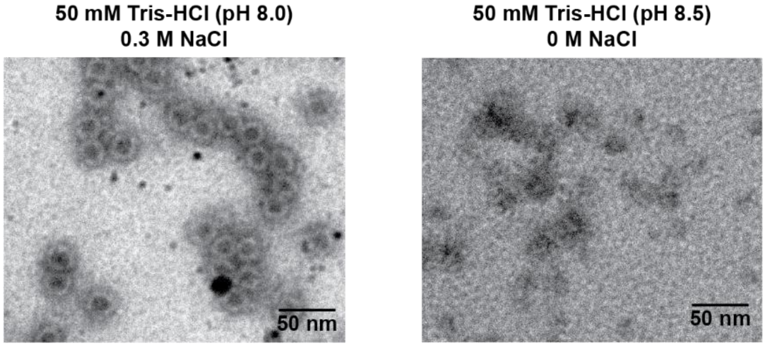


**Supplementary Figure 3. Salt-dependent assembly of cpAaLS(Strep).** Negative-stain TEM images of cpAaLS(strep) prepared in 50 mM Tris-HCl buffer (pH 8.0) containing 0.3 mM NaCl (left) or in 50 mM Tris-HCl buffer (pH 8.5) (right).

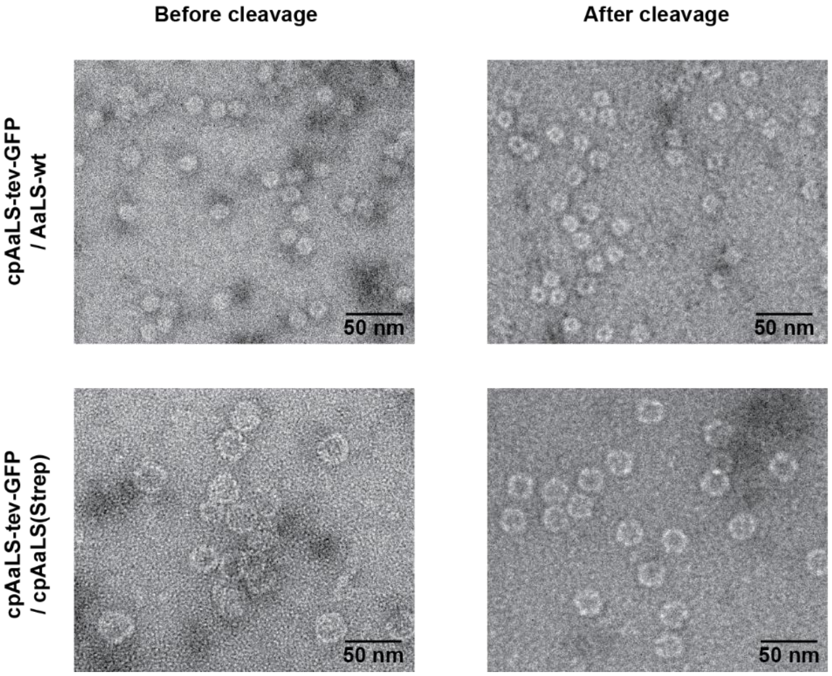


**Supplementary Figure 4. TEM images of AaLS cages containing cpAaLS-tev-GFP-SsrA.** These patchwork assemblies were obtained by coproduction of cpAaLS-tev-GFP-SsrA with either AaLS-wt (top) or cpAaLS(Strep) (bottom), followed by isolation using Strep-Tactin affinity chromatography. The TEM images were taken before (left) and after (right) treatment with TEV protease.

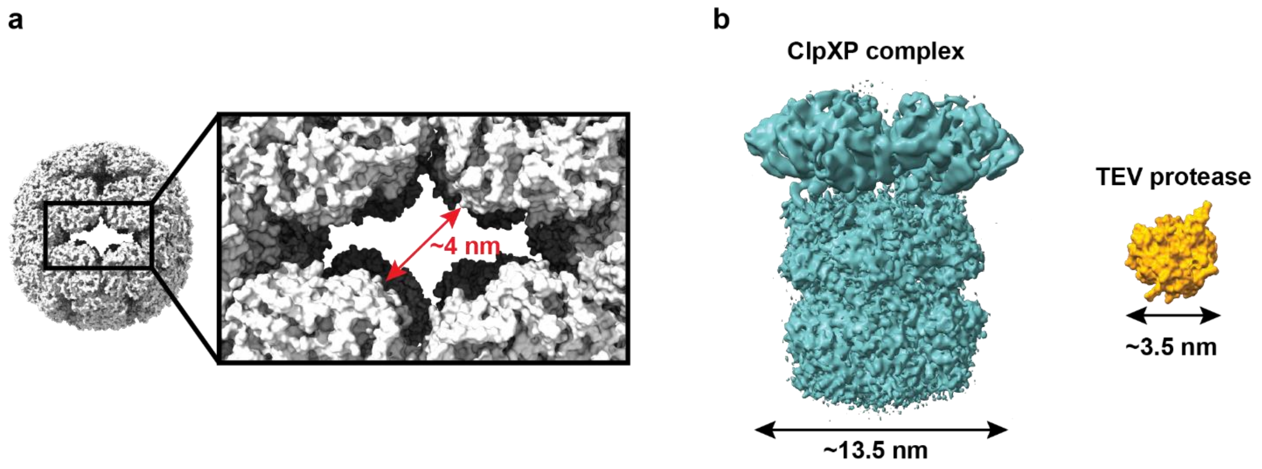


**Supplementary Figure 5. Size comparison between AaLS pores and proteases.** (a) CryoEM map of the cpAaLS 180mer cage (PDB: 9G3N) with an enlarged image of the pore region. (b) CryoEM map of ClpXP complex from *Listeria monocytogenes* (left, PDB: 6SFW) and crystal structure of TEV protease (right, PDB: 1LVB).

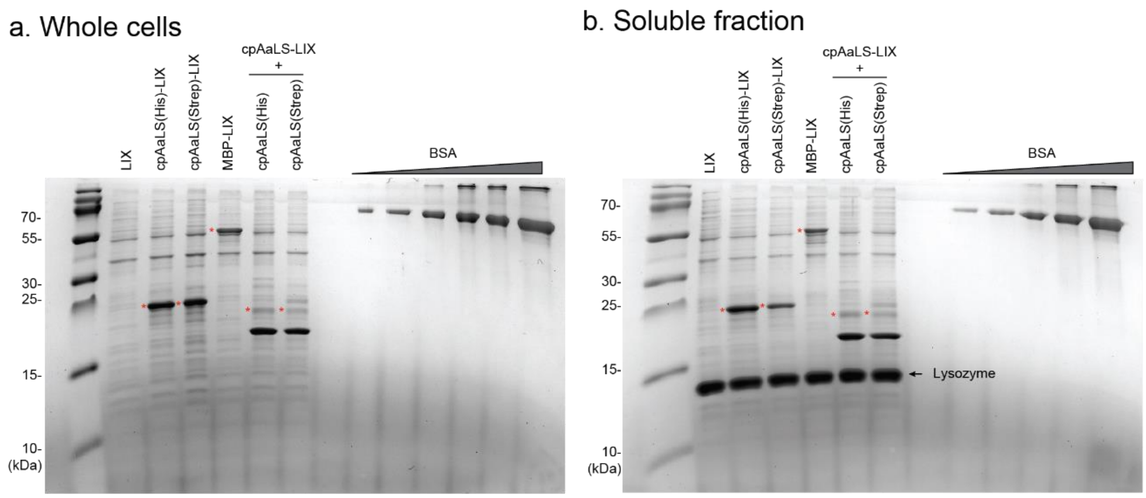


**Supplementary Figure 6. SDS-PAGE analysis of LIX production.** (a,b) LIX peptides were produced in different fusion designs in *E. coli* cells, and whole cells (a) and soluble fraction after cell lysis (b) were analyzed by SDS-PAGE. The bands containing LIX moieties are marked with red asterisks. Part of the gel images are also shown in Figure 4b. BSA (0.125-2 μg) samples were loaded on the same gels and used for quantification. Calculated molecular masses of proteins are; LIX, 6.6 kDa; cpAaLS(His)-LIX, 24.3 kDa; cpAaLS(Strep)-LIX, 24.6 kDa; MBP-LIX, 47.3 kDa; cpAaLS-LIX, 23.6 kDa. cpAaLS(His), 18.2 kDa; and cpAaLS(Strep), 18.5 kDa.

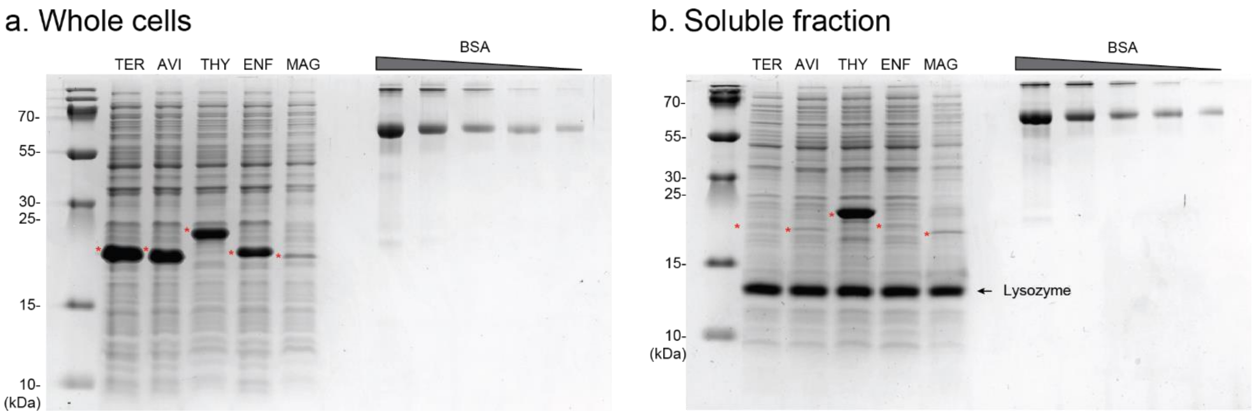


**Supplementary Figure 7. SDS-PAGE analysis of the therapeutic peptide production in single-component AaLS cages.** (a,b) TER, AVI, THY, ENF, and MAG peptides were produced as a fusion to cpAaLS(Strep) via TEV protease recognition sequence in *E. coli* cells, and whole cells (a) and soluble fraction after cell lysis (b) were analyzed by SDS-PAGE. The bands containing these peptide moieties (red asterisks) were used for densitometry analysis shown in Figure 4c. BSA (0.125-2 μg) samples were loaded on the same gels and used for quantification. Calculated molecular masses of proteins are; cpAaLS(Strep)-TER, 24.0 kDa; cpAaLS(Strep)-AVI, 23.1 kDa; cpAaLS(Strep)-THY, 19.8 kDa; cpAaLS(Strep)-ENF, 24.2 kDa; cpAaLS(Strep)-MAG, 22.2 kDa.

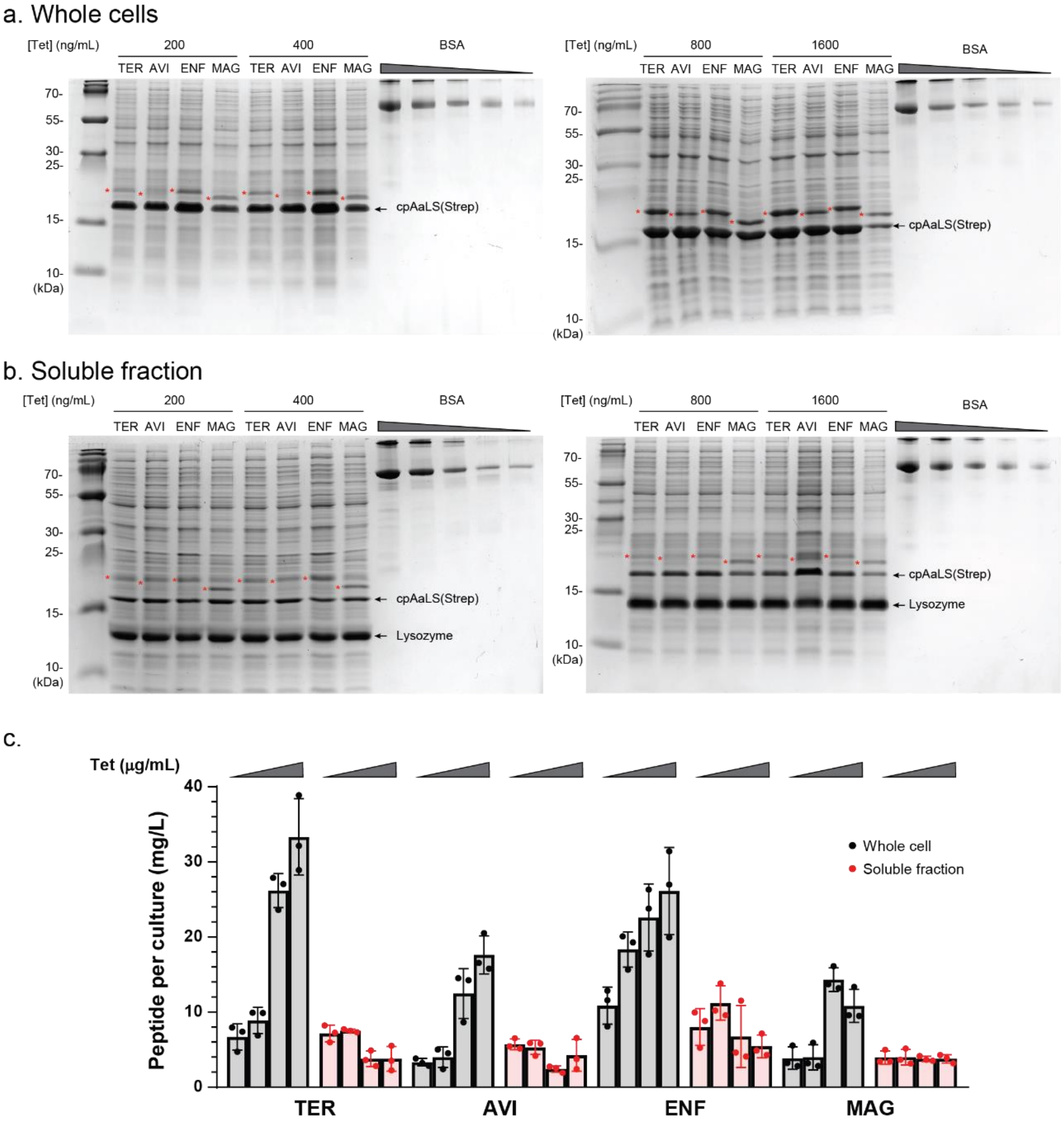


**Supplementary Figure 8. SDS-PAGE analysis of the therapeutic peptide production in patchwork AaLS cages.** (a,b) TER, AVI, THY, ENF, and MAG peptides were coproduced as a fusion to cpAaLS via TEV protease recognition sequence with cpAaLS(Strep) in *E. coli* cells, and whole cells (a) and soluble fraction after cell lysis (b) were analyzed by SDS-PAGE. The bands containing these peptide moieties (red asterisks) were used for densitometry analysis, as shown in panel (c). The results with 0.4 g/mL tetracycline are also shown in Figure 5a,c. BSA (0.125-2 µg) samples were loaded on the same gels and used for quantification. Data are shown as means ± standard deviation from three biological replicates.

Calculated molecular masses of proteins are; cpAaLS-TER, 22.8 kDa; cpAaLS-AVI, 22.0 kDa; cpAaLS-ENF, 23.1 kDa; cpAaLS-MAG, 21.2 kDa.

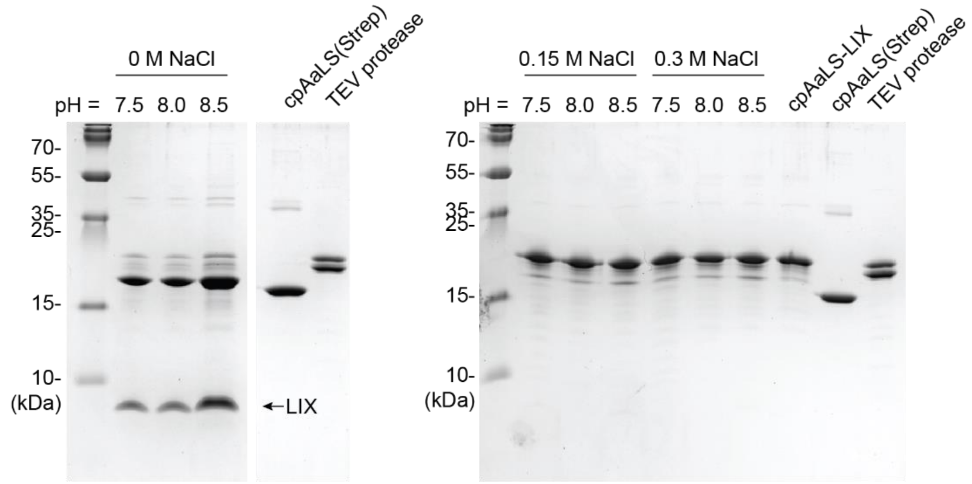


**Supplementary Figure 9. Salt-dependent cleavage of cpAaLS-LIX.** Part of the left gel is also shown in Figure 6a. The theoretical molecular masses of proteins are as follows: cpAaLS(Strep)-tev-LIX, 24.6 kDa; cpAaLS(Step), 18.5 kDa; TEVp, 28.8 kDa; LIX, 4.9 kDa.

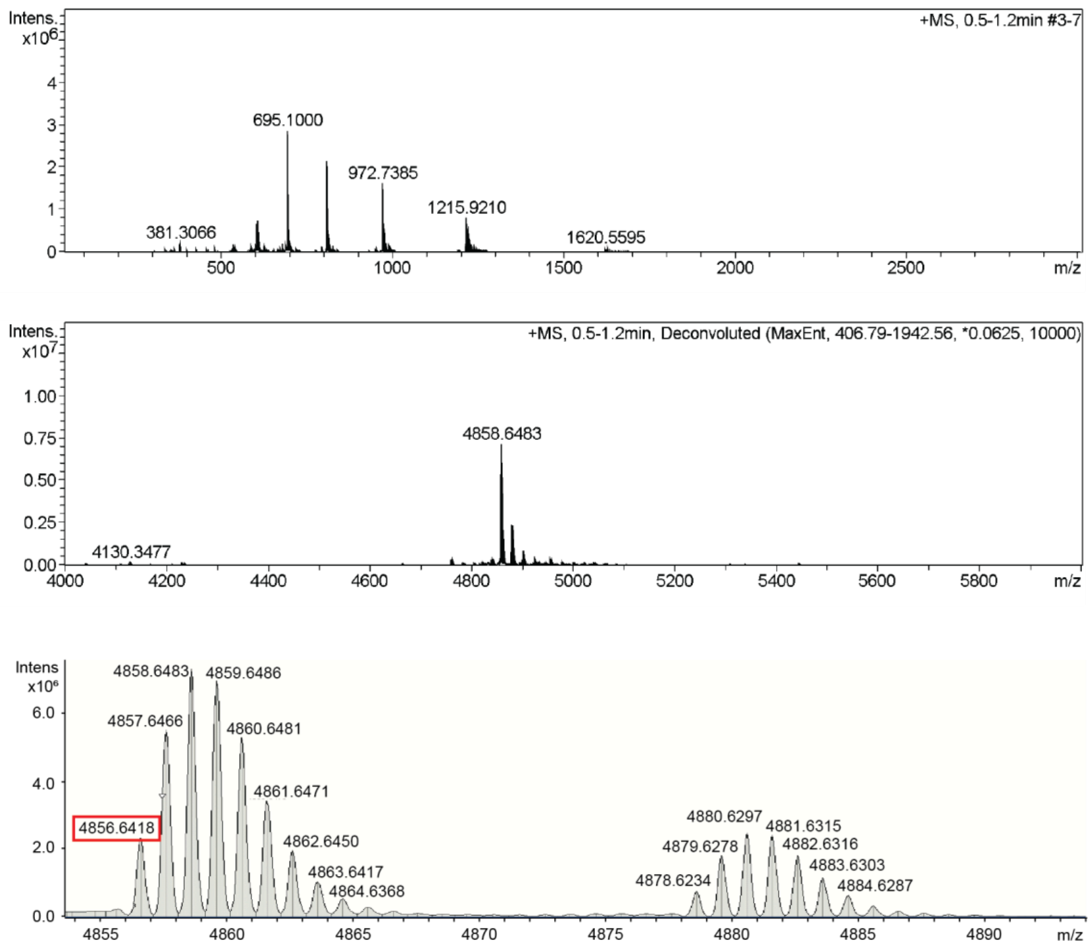


**Supplementary Figure 10. Mass spectrometry of LIX.** Electron spray ionization mass spectrum of purified LIX, shown as raw data (top), after peak deconvolution (middle), and enlarged profile of the main peaks. Part of the spectrum is also shown in Figure 6d. The calculated monoisotopic mass of LIX is 4856.53. The second-highest peak cluster is likely sodium adducts.

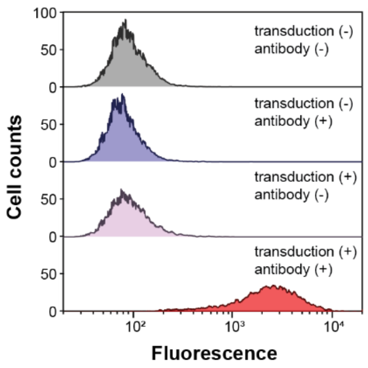


**Supplementary Figure 11. Flow cytometry analysis of the CRE/CREB luciferase reporter HEK cells producing GLP-1 receptor.** The cells were transduced with lentiviral vectors carrying a human GLP-1R gene [transduction (+)] and labeled with a human GLP-1R antibody conjugated with Alexa Fluor 488 [antibody (+)]. Those without the transduction [transduction (-)] and/or the antibody labeling [antibody (-)] are provided as controls. The dead cell population was excluded based on the propidium iodide (PI) fluorescence, and only live cells were analyzed for GLP-1R expression.

### Supplementary Tables

#### Supplementary Table 1. Amino acid sequence of the proteins used for this study

| Name | Sequence | M.W. | $\varepsilon_{280}$ |
|---|---|---|---|
| AaLS-wt | MEIYEGKLTAEGLRFGIVASRFNHALVDRLVEGAIDCIVRHGGREEDITLVR<br>VPGSWEIPVAAGELARKEDIDAVIAIGVLIRGATPHFDYIASEVSKGLANLS<br>LELRKPITFGVITADTLEQAIERAGTKHGNKGWEAALSAIEMANLFKSLRLE<br>GGWSHPQFEK* | 18,102 | 19,480 |
| cpAaLS (His) | MTLEQAIERAGTKHGNKGWEAALSAIEMANLFKSLRGTGHHHHHHGGSSMEI<br>YEGKLTAEGLRFGIVASRFNHALVDRLVEGAIDCIVRHGGREEDITLVRVPG<br>SWEIPVAAGELARKEDIDAVIAIGVLIRGATPHFDYIASEVSKGLANLSLEL<br>RKPITFGVITAD* | 18,164 | 13,980 |
| cpAaLS-GFP-SsrA | MTLEQAIERAGTKHGNKGWEAALSAIEMANLFKSLRGTGGSGSSMEIYEGKL<br>TAEGLRFGIVASRFNHALVDRLVEGAIDCIVRHGGREEDITLVRVPGSWEIP<br>VAAGELARKEDIDAVIAIGVLIRGATPHFDYIASEVSKGLANLSLELRKPIT<br>FGVITADAGGAGGSGMASKGEELFTGVVPILVELDGDVNGHKFSVSGEGEGD<br>ATYGKLTLKFICTTGKLPVPWPTLVTTLCYGVQCFSRYPDHMKRHDFFKSAM<br>PEGYVQERTIFFKDDGNYKTRAEVKFEGDTLVNRIELKGIDFKEDGNILGHK<br>LEYNYNSHNVYIMADKQKNGIKVNFKTRHNIEDGSVQLADHYQQNTPIGDGP<br>VLLPDNHYLSTQSALSKDPNEKRDHMVLLEFVTAAGITHGMDELYKSGGSMA<br>LEAANDENYALAA* | 46,717 | - |
| cpAaLS-tev*-GFP-ssrA | MTLEQAIERAGTKHGNKGWEAALSAIEMANLFKSLRGTGGSGSSMEIYEGKL<br>TAEGLRFGIVASRFNHALVDRLVEGAIDCIVRHGGREEDITLVRVPGSWEIP<br>VAAGELARKEDIDAVIAIGVLIRGATPHFDYIASEVSKGLANLSLELRKPIT<br>FGVITADAGGAENLYFQSGGSGMASKGEELFTGVVPILVELDGDVNGHKFSV<br>SGEGEGDATYGKLTLKFICTTGKLPVPWPTLVTTLCYGVQCFSRYPDHMKRH<br>DFFKSAMPEGYVQERTIFFKDDGNYKTRAEVKFEGDTLVNRIELKGIDFKED<br>GNILGHKLEYNYNSHNVYIMADKQKNGIKVNFKTRHNIEDGSVQLADHYQQN<br>TPIGDGPVLLPDNHYLSTQSALSKDPNEKRDHMVLLEFVTAAGITHGMDELY<br>KSGGSMALEAANDENYALAA* | 47,599 | - |
| cpAaLS (Strep) | MTLEQAIERAGTKHGNKGWEAALSAIEMANLFKSLRGTGGWSHPQFEKGTSS<br>MEIYEGKLTAEGLRFGIVASRFNHALVDRLVEGAIDCIVRHGGREEDITLVR<br>VPGSWEIPVAAGELARKEDIDAVIAIGVLIRGATPHFDYIASEVSKGLANLS<br>LELRKPITFGVITAD* | 18,482 | 19,480 |
| cpAaLS (Strep)-tev*-LIX | MTLEQAIERAGTKHGNKGWEAALSAIEMANLFKSLRGTGGWSHPQFEKGTSS<br>MEIYEGKLTAEGLRFGIVASRFNHALVDRLVEGAIDCIVRHGGREEDITLVR<br>VPGSWEIPVAAGELARKEDIDAVIAIGVLIRGATPHFDYIASEVSKGLANLS<br>LELRKPITFGVITADAGGAGGSGENLYFQHGEGTFTSDLSKQMEEEAVRLFI<br>EWLKNGGPSSGAPPSKKKKKK* | 24,633 | 26,470 |
| cpAaLS-tev*-LIX | MTLEQAIERAGTKHGNKGWEAALSAIEMANLFKSLRGTGGSGSSMEIYEGKL<br>TAEGLRFGIVASRFNHALVDRLVEGAIDCIVRHGGREEDITLVRVPGSWEIP<br>VAAGELARKEDIDAVIAIGVLIRGATPHFDYIASEVSKGLANLSLELRKPIT<br>FGVITADAGGAGGSGENLYFQHGEGTFTSDLSKQMEEEAVRLFIEWLKNGGP<br>SSGAPPSKKKKKK* | 23,579 | 20,970 |
| Mal-LIX | MKIHHHHHHEEGKLVIWINGDKGYNGLAEVGKKFEKDTGIKVTVEHPDKLEE<br>KFPQVAATGDGPDIIFWAHDRFGGYAQSGLLAEITPDKAFQDKLYPFTWDAV<br>RYNGKLIAYPIAVEALSLIYNKDLLPNPPKTWEEIPALDKELKAKGKSALMF<br>NLQEPYFTWPLIAADGGYAFKYENGKYDIKDVGVDNAGAKAGLTFLVDLIKN<br>KHMNADTDYSIAEAAFNKGETAMTINGPWAWSNIDTSKVNYGVTVLPTFKGQ<br>PSKPFVGVLSAGINAASPNKELAKEFLENYLLTDEGLEAVNKDKPLGAVALK<br>SYEEELAKDPRIAATMENAQKGEIMPNIPQMSAFWYAVRTAVINAASGRQTV<br>DEALKDAQTAGGAGGSGENLYFQHGEGTFTSDLSKQMEEEAVRLFIEWLKNG<br>GPSSGAPPSKKKKKK* | 47,314 | 73,340 |

| | | | |
|---|---|---|---|
| LIX | MHHHHHHENLYFQHGEGTFTSDLSKQMEEEAVRLFIEWLKNGGPSSGAPPSKKKKKK* | 6,608 | 6,990 |
| cpAaLS (Strep)-tev*-TER | MTLEQAIERAGTKHGNKGWEAALSAIEMANLFKSLRGTGGWSHPQFEKGTSSMEIYEGKLTAEGLRFGIVASRFNHALVDRLVEGAIDCIVRHGGREEDITLVRVPGSWEIPVAAGELARKEDIDAVIAIGVLIRGATPHFDYIASEVSKGLANLSLELRKPITFGVITADAGGAGGSGENLYFQSVSEIQLMHNLGKHLNSMERVEWLRKKLQDVHNF* | 23,891 | 26,470 |
| cpAaLS (Strep)-tev*-AVI | MTLEQAIERAGTKHGNKGWEAALSAIEMANLFKSLRGTGGWSHPQFEKGTSSMEIYEGKLTAEGLRFGIVASRFNHALVDRLVEGAIDCIVRHGGREEDITLVRVPGSWEIPVAAGELARKEDIDAVIAIGVLIRGATPHFDYIASEVSKGLANLSLELRKPITFGVITADAGGAGGSGENLYFQHSDAVFTDNYTRLRKQMAVKKYLNSILN* | 23,100 | 23,950 |
| cpAaLS (Strep)-tev*-THY | MTLEQAIERAGTKHGNKGWEAALSAIEMANLFKSLRGTGGWSHPQFEKGTSSMEIYEGKLTAEGLRFGIVASRFNHALVDRLVEGAIDCIVRHGGREEDITLVRVPGSWEIPVAAGELARKEDIDAVIAIGVLIRGATPHFDYIASEVSKGLANLSLELRKPITFGVITADAGGAGGSGENLYFQ* | 19,791 | 20,970 |
| cpAaLS (Strep)-tev*-ENF | MTLEQAIERAGTKHGNKGWEAALSAIEMANLFKSLRGTGGWSHPQFEKGTSSMEIYEGKLTAEGLRFGIVASRFNHALVDRLVEGAIDCIVRHGGREEDITLVRVPGSWEIPVAAGELARKEDIDAVIAIGVLIRGATPHFDYIASEVSKGLANLSLELRKPITFGVITADAGGAGGSGENLYFQYTSLIHSLIEESQNQQEKNEQELLELDKWASLWNWF* | 24,224 | 38,960 |
| cpAaLS (Strep)-tev*-MAG | MTLEQAIERAGTKHGNKGWEAALSAIEMANLFKSLRGTGGWSHPQFEKGTSSMEIYEGKLTAEGLRFGIVASRFNHALVDRLVEGAIDCIVRHGGREEDITLVRVPGSWEIPVAAGELARKEDIDAVIAIGVLIRGATPHFDYIASEVSKGLANLSLELRKPITFGVITADAGGAGGSGENLYFQGIGKFLHSAKKFGKAFVGEIMNS* | 22,240 | 20,970 |
| cpAaLS-tev*-TER | MTLEQAIERAGTKHGNKGWEAALSAIEMANLFKSLRGTGGSGSSMEIYEGKLTAEGLRFGIVASRFNHALVDRLVEGAIDCIVRHGGREEDITLVRVPGSWEIPVAAGELARKEDIDAVIAIGVLIRGATPHFDYIASEVSKGLANLSLELRKPITFGVITADAGGAGGSGENLYFQSVSEIQLMHNLGKHLNSMERVEWLRKKLQDVHNF* | 22,837 | 20,970 |
| cpAaLS-tev*-AVI | MTLEQAIERAGTKHGNKGWEAALSAIEMANLFKSLRGTGGSGSSMEIYEGKLTAEGLRFGIVASRFNHALVDRLVEGAIDCIVRHGGREEDITLVRVPGSWEIPVAAGELARKEDIDAVIAIGVLIRGATPHFDYIASEVSKGLANLSLELRKPITFGVITADAGGAGGSGENLYFQHSDAVFTDNYTRLRKQMAVKKYLNSILN* | 22,046 | 18,450 |
| cpAaLS-tev*-ENF | MTLEQAIERAGTKHGNKGWEAALSAIEMANLFKSLRGTGGSGSSMEIYEGKLTAEGLRFGIVASRFNHALVDRLVEGAIDCIVRHGGREEDITLVRVPGSWEIPVAAGELARKEDIDAVIAIGVLIRGATPHFDYIASEVSKGLANLSLELRKPITFGVITADAGGAGGSGENLYFQYTSLIHSLIEESQNQQEKNEQELLELDKWASLWNWF* | 23,170 | 33,460 |
| cpAaLS-tev*-MAG | MTLEQAIERAGTKHGNKGWEAALSAIEMANLFKSLRGTGGSGSSMEIYEGKLTAEGLRFGIVASRFNHALVDRLVEGAIDCIVRHGGREEDITLVRVPGSWEIPVAAGELARKEDIDAVIAIGVLIRGATPHFDYIASEVSKGLANLSLELRKPITFGVITADAGGAGGSGENLYFQGIGKFLHSAKKFGKAFVGEIMNS* | 21,186 | 15,470 |
| Strep-sTEVp-R5 | GWSHPQFEKGGSGESLFKGPRDYNPISSTIVHLTNESDGHTTSLYGIGFGPFIITNKHLFRRNNGTLVVQSLHGVFKVKNTTTLQQHLIDGRDMIIIRMPKDFPPFPQKLKFREPQREERIVLVTTNFQTKSMSSMVSDTSSTFPSGDGIFWKHWIQTKDGQCGSPLVSTRDGFIVGIHSASNFTNTNNYFTSVPKNFMELLTNQEAQQWVSGWRLNADSVLWGGHKVFMDKPEEPFQPVKEATQLMNRRRRR* | 28,846 | 37,470 |

**Supplementary Table 2. Therapeutic peptides produced in this study**

| Peptide | #AA[a] | Sequence[b] | Biological activity | Hydrophobic AA (%)[c] | GRAVY[d] |
|---|---|---|---|---|---|
| Lixisenatide (LIX) | 44 | **HGEGTFTSDLSKQMEE EAVRLFIEWLKNGGPS SGAPPSKKKKKK** | glucagon-like peptide-1 receptor (GLP1R) agonist | 31.8 | -1.11 |
| Teriparatide (TER) | 34 | **SVSEIQLMHNLGKHLN SMERVEWLRKKLQDVH NF** | parathyroid hormone (PTH) | 38.2 | -0.68 |
| Aviptadil (AVI) | 28 | **HSDAVFTDNYTRLRKQ MAVKKYLNSILN** | melanocortin 1 receptor agonist | 35.2 | -0.64 |
| Thymosin beta 4 (THY) | 43 | **SDKPDMAEIEKFDKSK LKKTETQEKNPLPSKE TIEQEKQAGES** | VPAC 1 & 2 agonist | 25.6 | -1.70 |
| Enfuvirtide (ENF) | 36 | **YTSLIHSLIEESQNQQ EKNEQELLELDKWASL WNWF** | HIV-1 gp41 fusion inhibitor | 36.1 | -0.88 |
| Magainin 2 (MAG) | 23 | **GIGKFLHSAKKFGKAF VGEIMNS** | antimicrobial peptide | 43.4 | 0.083 |

a. The number of amino acids.

b. The N- and C-termini of these peptides produced in this study remain free amine and carboxylate.

c. Calculated by peptide 2.0 (https://www.peptide2.com/N_peptide_hydrophobicity_hydrophilicity.php).

d. Grand average of hydropathicity, calculated with ProtParam (https://web.expasy.org/protparam/).

**Supplementary Table 3. Plasmids used for this study**

| name | gene | N-tag | C-tag | promoter /operator[a] | Ori | marker[b] | ref |
|---|---|---|---|---|---|---|---|
| pET15_AaLS-wt-Strep | lumazine synthase from *Aquifix aeolicus* (AaLS) | - | StrepII | $P_{T7}$ / *lac*O | ColE1 | $Amp^R$ | This study |
| pET15_cpAaLS(His) | circularly permuted variant of AaLS (cpAaLS(119)) | 6x His (loop) | | $P_{T7}$ / *lac*O | ColE1 | $Amp^R$ | This study |
| pMG_cpAaLS (Strep) | circularly permuted variant of AaLS (cpAaLS(119)) | Strep II (loop) | | $P_{T7}$ / *lac*O and $P_{sal}$ | pBR322 | $Amp^R$ | 20 |
| pET15_cpAaLS(Strep) | cpAaLS(119) | Strep II (loop) | | $P_{T7}$ / *lac*O | ColE1 | $Amp^R$ | This study |
| pACYC_Ptet_ cpAaLS-GFP-SsrA | red-shifted GFP (rsGFP) fused to cpAaLS(119) | - | SsrA | $P_{tet}$ / *tet*O | p15A | $Cm^R$ | 20 |
| pACYC_Ptet_ cpAaLS-GFP | rsGFP fused to cpAaLS(119) | - | - | $P_{tet}$ / *tet*O | p15A | $Cm^R$ | 20 |
| pACYC_Ptet_H-GFP-SsrA | rsGFP | 6x His | SsrA | $P_{tet}$ / *tet*O | p15A | $Cm^R$ | 20 |
| pACYC_Ptet_H-GFP | rsGFP | 6x His | - | $P_{tet}$ / *tet*O | p15A | $Cm^R$ | 20 |
| pACYC_Ptet_ cpAaLS-tev*-GFP-ssrA | rsGFP fused to cpAaLS(119) via TEV protease recognition site | - | SsrA | $P_{tet}$ / *tet*O | p15A | $Cm^R$ | This study |
| pET15_cpAaLS(Strep)-tev*-LIX | Lixisenatide (LIX) fused to cpAaLS(119) via TEV protease recognition site | Strep II (loop) | | $P_{T7}$ / *lac*O | ColE1 | $Amp^R$ | This study |
| pACYC_Ptet_cpAaLS-tev*-LIX | Lixisenatide (LIX) fused to cpAaLS(119) via TEV protease recognition site | - | - | $P_{tet}$ / *tet*O | p15A | $Cm^R$ | This study |

| | | | | | | | |
|---|---|---|---|---|---|---|---|
| pET15_H-MBP-tev*-LIX | Lixisenatide (LIX) fused to maltose-binding protein via TEV protease recognition site | 6x His | - | $P_{T7}$ / *lac*O | ColE1 | $Amp^R$ | This study |
| pET15_H-tev*-LIX | Lixisenatide (LIX) fused TEV protease recognition site | 6x His | - | $P_{T7}$ / *lac*O | ColE1 | $Amp^R$ | This study |
| pET15_cpAaLS(Strep)-tev*-TER | Teriparatide (TER) fused to cpAaLS(119) via TEV protease recognition site | Strep II (loop) | | $P_{T7}$ / *lac*O | ColE1 | $Amp^R$ | This study |
| pET15_cpAaLS(Strep)-tev*-AVI | Aviptadil (LIX) fused to cpAaLS(119) via TEV protease recognition site | Strep II (loop) | | $P_{T7}$ / *lac*O | ColE1 | $Amp^R$ | This study |
| pET15_cpAaLS(Strep)-tev*-THY | thymosin beta-4 (THY) fused to cpAaLS(119) via TEV protease recognition site | Strep II (loop) | | $P_{T7}$ / *lac*O | ColE1 | $Amp^R$ | This study |
| pET15_cpAaLS(Strep)-tev*-ENF | Enfuvirtide (ENF) fused to cpAaLS(119) via TEV protease recognition site | Strep II (loop) | | $P_{T7}$ / *lac*O | ColE1 | $Amp^R$ | This study |
| pET15_cpAaLS(Strep)-tev*-MAG | Magainine 2 (MAG) fused to cpAaLS(119) via TEV protease recognition site | Strep II (loop) | | $P_{T7}$ / *lac*O | ColE1 | $Amp^R$ | This study |
| pACYC_Ptet_cpAaLS-tev*-TER | Teriparatide (TER) fused to cpAaLS(119) via TEV protease recognition site | - | - | $P_{tet}$ / *tet*O | p15A | $Cm^R$ | This study |
| pACYC_Ptet_cpAaLS-tev*-AVI | Aviptadil (LIX) fused to cpAaLS(119) via TEV protease recognition site | - | - | $P_{tet}$ / *tet*O | p15A | $Cm^R$ | This study |
| pACYC_Ptet_cpAaLS-tev*-ENF | Enfuvirtide (ENF) fused to cpAaLS(119) via TEV | - | - | $P_{tet}$ / *tet*O | p15A | $Cm^R$ | This study |

| | | | | | | | |
|---|---|---|---|---|---|---|---|
| | protease recognition site | | | | | | |
| pACYC_Ptet_cpAaLS-tev*-MAG | Magainine 2 (MAG) fused to cpAaLS(119) via TEV protease recognition site | - | - | $P_{tet}$ / *tet*O | p15A | $Cm^R$ | This study |
| pMAL_S-sTEVp | engineered variant of protease from tobacco etch virus (super TEV protease) | Strep II | - | $P_{tac}$ / *lac*O | ColE1 | $Amp^R$ | This study |

a. $P_{T7}$/*lacO,* T7 promoter combined with lactose operator and the LacI repressor (*lacI*); $P_{sal}$, the salicylate promoter regulated by the NahR transcriptional activator (*nahR*); $P_{tet}$/*tetO*, the tetracycline promoter combined with tetracycline operator and the TetR repressor (*tetR*); $P_{tac}$, tryptophan-lactose UV5 (tac) promoter

b. $Amp^R$, β-lactamase; $Cm^R$, chloramphenicol acetyltransferase

#### Supplementary Table 4. Oligonucleotides used for this study

| name | sequence |
|---|---|
| FW_BglI_TEV_BamHI | GGGCGGAAAATCTGTATTTTCAGAGCGGCG |
| RV_BamHI_TEV_BglI | GATCCGCCGCTCTGAAAATACAGATTTTCCGCCCCAC |
| FW_pACYC184 | GGATCTGCATCGCAGGA |
| RV_pACYC184 | AAGCTTATCGATGATAAGCTGT |
| TetR_Ptet_cpAaLS-MAG2 | GTTGTAATTCTCATGTTTGACAGCTTATCATCGATAAGCTTGCATGCTTAAGACCCACTTTCACATTTAAGTTGTTTTTCTAATCCGCAAATGATCAATTCAAGGCCGAATAAGAAGGCTGGCTCTGCACCTTGGTGATCAAATAATTCGATAGCTTGTCGTAATAATGGCGGCATACTATCAGTAGTAGGTGTTTCCCTTTCTTCTTTAGCGACTTGATGCTCTTGATCTTCCAATACGCAACCTAAAGTAAAATGCCCCACAGCGCTGAGTGCATATAATGCATTCTCTAGTGAAAAACCTTGTTGGCATAAAAAGGCTAATTGATTTTCGAGAGTTTCATACTGTTTTTCTGTAGGCCGTGTACCTAAATGTACTTTTGCTCCATCGCGATGACTTAGTAAAGCACATCTAAAACTTTTAGCGTTATTACGTAAAAAATCTTGCCAGCTTTCCCCTTCTAAAGGGCAAAAGTGAGTATGGTGCCTATCTAACATCTCAATGGCTAAGGCGTCGAGCAAAGCCCGCTTATTTTTTACATGCCAATACAATGTAGGCTGCTCTACACCTAGCTTCTGGGCGAGTTTACGGGTTGTTAAACCTTCGATTCCGACCTCATTAAGCAGCTCTAATGCGCTGTTAATCACTTTACTTTTATCTAATCTCGACATCATTAATTCCTAATTTTTGTTGACACTCTATCATTGATAGAGTTATTTTACCACTCCCTATCAGTGATAGAGAAAAGTCTAGAAATAATTTTGTTTAACTTTAAGAAGGAGATATACATATGACCTTGGAACAGGCTATCGAGCGCGCCGGCACAAAACACGGCAACAAAGGTTGGGAAGCAGCGCTTTCTGCCATTGAAATGGCAAACTTATTCAAGTCTCTCCGAGGTACCGGTGGCTCGGGGAGCTCGATGGAAATCTACGAAGGTAAACTAACTGCTGAAGGCCTTCGTTTCGGTATCGTAGCATCACGTTTTAATCATGCTCTTGTCGACCGTCTGGTGGAGGGTGCAATTGATTGCATAGTCCGTCATGGCGGCCGTGAAGAAGACATTACTCTGGTTCGTGTTCCAGGCTCATGGGAAATACCGGTTGCTGCGGGTGAACTGGCGCGTAAAGAGGACATTGATGCTGTTATCGCAATTGGCGTTCTCATCAGAGGCGCAACGCCACATTTCGATTATATCGCCTCTGAAGTTTCAAAAGGCCTCGCGAACCTTTCATTAGAACTACGTAAACCTATCACCTTCGGTGTTATTACAGCTGACGCCGGTGGGGCGGGCGGATCCGGTGAAAATCTGTATTTTCAGAGCGGTATTGGTAAATTTCTGCATAGCGCGAAAAAATTTGGTAAAGCGTTTGTGGGTGAAATTATGAACAGCTAATCTAGTCAGCTGATCCGGCTGCTAACAAAGCCCGAAAGGAAGCTGAGTTGGCTGCTGCCACCGCTGAGCAATAACTAGCATAACCCCTTGGGGCCTCTAAACGGGTCTTGAGGGGTTTTTTGCTGAAAGGAGGAACTATATCCGGATGCGGATCTGCATCGCAGGATGCTGCTGGCTACCCTGTGGAAC |
| FW_uni | TAATCTAGTCAGCTGATCCG |
| RV_uni | GCTCTGAAAATACAGATTTTCAC |
| FW_LIX_1 | AAAATCTGTATTTTCAGAGCCATGGGGAGGGCACGTTTACGAGTGATCTTTCCAAGCAAATGGAGGAAGAGGCAGTTCGGCTGTTTATCG |
| RV_LIX_1 | CGATAAACAGCCGAACTGCCTCTTCCTCCATTTGCTTGGAAAGATCACTCGTAAACGTGCCCTCCCCATGGCTCTGAAAATACAGATTTT |
| FW_LIX_2 | GGCAGTTCGGCTGTTTATCGAGTGGTTGAAAAATGGTGGCCCTTCGAGCGGCGCTCCTCCTAGCAAGAAAAAAAAAAGAAATAATCTAGTCAGCTGATCCG |
| RV_LIX_2 | CGGATCAGCTGACTAGATTATTTCTTTTTTTTTTTCTTGCTAGGAGGAGCGCCGCTCGAAGGGCCACCATTTTTCAACCACTCGATAAACAGCCGAACTGCC |
| FW_TER_1 | AAAATCTGTATTTTCAGAGCAGTGTGAGCGAAATCCAACTTATGCATAATCTCGGGAAACATTTAAATTCAATGGAGCGG |
| RV_TER_1 | CCGCTCCATTGAATTTAAATGTTTCCCGAGATTATGCATAAGTTGGATTTCGCTCACACTGCTCTGAAAATACAGATTTT |
| FW_TER_2 | ATTTAAATTCAATGGAGCGGGTCGAGTGGCTTCGTAAGAAGTTACAGGATGTACATAACTTCTAATCTAGTCAGCTGATCCG |
| RV_TER_2 | CGGATCAGCTGACTAGATTAGAAGTTATGTACATCCTGTAACTTCTTACGAAGCCACTCGACCCGCTCCATTGAATTTAAAT |
| FW_AVI_1 | AAAATCTGTATTTTCAGAGCCACTCAGACGCCGTGTTTACTGACAACTATACTCGGCTTCGTAAACAAAT |
| RV_AVI_1 | ATTTGTTTACGAAGCCGAGTATAGTTGTCAGTAAACACGGCGTCTGAGTGGCTCTGAAAATACAGATTTT |
| FW_AVI_2 | ACTCGGCTTCGTAAACAAATGGCAGTTAAAAAATATCTCAACTCCATTCTTAATTAATCTAGTCAGCTGATCCG |
| RV_AVI_2 | CGGATCAGCTGACTAGATTAATTAAGAATGGAGTTGAGATATTTTTTAACTGCCATTTGTTTACGAAGCCGAGT |
| FW_ENF_1 | AAAATCTGTATTTTCAGAGCTACACGAGTTTGATTCACAGTCTTATTGAGGAAAGCCAAAATCAACAAGAGAAAAACGAG |
| RV_ENF_1 | CTCGTTTTTCTCTTGTTGATTTTGGCTTTCCTCAATAAGACTGTGAATCAAACTCGTGTAGCTCTGAAAATACAGATTTT |
| FW_ENF_2 | ATCAACAAGAGAAAAACGAGCAAGAGTTGTTGGAGCTTGACAAATGGGCATCGCTTTGGAATTGGTTCTAATCTAGTCAGCTGATCCG |

| | |
|---|---|
| RV_ENF_2 | CGGATCAGCTGACTAGATTAGAACCAATTCCAAAGCGATGCCCATTTGTCAAGCTCCAACAACTCTTGCTCGTTT<br>TTCTCTTGTTGAT |
| FW_THY_1 | AAAATCTGTATTTTCAGAGCTCTGACAAGCCGGATATGGCGGAGATTGAAAAGTTTGATAAGTCTAAACTTAAAA<br>AGACAGAGACTCAAGAAAAAAACCC |
| RV_THY_1 | GGGTTTTTTTCTTGAGTCTCTGTCTTTTTAAGTTTAGACTTATCAAACTTTTCAATCTCCGCCATATCCGGCTTG<br>TCAGAGCTCTGAAAATACAGATTTT |
| FW_THY_2 | GAGACTCAAGAAAAAAACCCATTACCGTCTAAAGAGACAATCGAACAGGAAAAACAGGCGGGGGAGTCGTAATCT<br>AGTCAGCTGATCCG |
| RV_THY_2 | CGGATCAGCTGACTAGATTACGACTCCCCCGCCTGTTTTTCCTGTTCGATTGTCTCTTTAGACGGTAATGGGTTT<br>TTTTCTTGAGTCTC |
| FW_pET15b | TAACAAAGCCCGAAAGGAAGCTGAG |
| RV_pET15b | GTCAGCTGTAATAACACCGAAGGTGATAG |
| FW_MAG | CTTCGGTGTTATTACAGCTGACGC |
| RV_MAG | CAGCTTCCTTTCGGGCTTTGTTA |
| FW_Strep16_ins | CGGTGGGTGGAGCCATCCGCAGTTCGAAAAGGGGACGAGCT |
| RV_Strep16_ins | CGTCCCCTTTTCGAACTGCGGATGGCTCCACCCACCGGTAC |
| FW_del-LIX | GTGAAAATCTGTATTTTCAGCATGGGGAGGGCACG |
| RV_del-LIX | CGTGCCCTCCCCATGCTGAAAATACAGATTTTCAC |
| FW_del-TER | GGTGAAAATCTGTATTTTCAGAGTGTGAGCGAAATCCAACT |
| RV_del-TER | AGTTGGATTTCGCTCACACTCTGAAAATACAGATTTTCACC |
| FW_del-AVI | GGTGAAAATCTGTATTTTCAGCACTCAGACGCCGTGT |
| RV_del-AVI | ACACGGCGTCTGAGTGCTGAAAATACAGATTTTCACC |
| FW_del-THY | CCGGTGAAAATCTGTATTTTCAGTCTGACAAGCCGGATAT |
| RV_del-THY | ATATCCGGCTTGTCAGACTGAAAATACAGATTTTCACCGG |
| FW_del-ENF | CGGTGAAAATCTGTATTTTCAGTACACGAGTTTGATTCACAGTC |
| RV_del-ENF | GACTGTGAATCAAACTCGTGTACTGAAAATACAGATTTTCACCG |
| FW_del-MAG | GGTGAAAATCTGTATTTTCAGGGTATTGGTAAATTTCTGCA |
| FW_del-MAG | TGCAGAAATTTACCAATACCCTGAAAATACAGATTTTCACC |
| MalE | ATGAAAATCCACCATCACCATCATCACGAAGAAGGTAAACTGGTAATCTGGATTAACGGCGATAAAGGCTATAAC<br>GGTCTCGCTGAAGTCGGTAAGAAATTCGAGAAAGATACCGGAATTAAAGTCACCGTTGAGCATCCGGATAAACTG<br>GAAGAGAAATTCCCACAGGTTGCGGCAACTGGCGATGGCCCTGACATTATCTTCTGGGCACACGACCGCTTTGGT<br>GGCTACGCTCAATCTGGCCTGTTGGCTGAAATCACCCCGGACAAAGCGTTCCAGGACAAGCTGTATCCGTTTACC<br>TGGGATGCCGTACGTTACAACGGCAAGCTGATTGCTTACCCGATCGCTGTTGAAGCGTTATCGCTGATTTATAAC<br>AAAGATCTGCTGCCGAACCCGCCAAAAACCTGGGAAGAGATCCCGGCGCTGGATAAAGAACTGAAAGCGAAAGGT<br>AAGAGCGCGCTGATGTTCAACCTGCAAGAACCGTACTTCACCTGGCCGCTGATTGCTGCTGACGGGGGTTATGCG<br>TTCAAGTATGAAAACGGCAAGTACGACATTAAAGACGTGGGCGTGGATAACGCTGGCGCGAAAGCGGGTCTGACC<br>TTCCTGGTTGACCTGATTAAAAACAAACACATGAATGCAGACACCGATTACTCCATCGCAGAAGCTGCCTTTAAT<br>AAAGGCGAAACAGCGATGACCATCAACGGCCCGTGGGCATGGTCCAACATCGACACCAGCAAAGTGAATTATGGT<br>GTAACGGTACTGCCGACCTTCAAGGGTCAACCATCCAAACCGTTCGTTGGCGTGCTGAGCGCAGGTATTAACGCC<br>GCCAGTCCGAACAAAGAGCTGGCAAAAGAGTTCCTCGAAAACTATCTGCTGACTGATGAAGGTCTGGAAGCGGTT<br>AATAAAGACAAACCGCTGGGTGCCGTAGCGCTGAAGTCTTACGAGGAAGAGTTGGCGAAAGATCCACGTATTGCC<br>GCCACCATGGAAAACGCCCAGAAAGGTGAAATCATGCCGAACATCCCGCAGATGTCCGCTTTCTGGTATGCCGTG<br>CGTACTGCGGTGATCAACGCCGCCAGCGGTCGTCAGACTGTCGATGAAGCCCTGAAAGACGCGCAGACT |
| FW_pET15b_2 | CCCTGAAAGACGCGCAGACTGCCGGTGGGGCGGGCG |
| RV_pET15b_2 | TGGTGATGGTGGATTTTCATGGTATATCTCCTTCTTAAAGTTAAACAA |
| FW_uni_2 | TAACAAAGCCCGAAAGGAAG |
| RV_uni_2 | GCTCTGAAAATACAGATTTTCAC |
| LIX_FW_3 | GGTGAAAATCTGTATTTTCAGAGC |
| LIX_RV_3 | CTTCCTTTCGGGCTTTGTTA |
| FW_His-LIX | GTGGTGGTGATGATGGTGCATGGTATATCTCCTTCTTAAAGT |
| RV_His-LIX | ATGCACCATCATCACCACCACGAAAATCTGTATTTTCAGCATGGG |
| FW_BamHI_sTEVp | GAGCGGATCCGGTGAAAGCCTGTTTAAAGGTCC |
| RV_rrnB-T1-term | CAGTCTTTCGACTGAGCCTTTCG |

### Detailed experimental procedures

#### Materials

HisPur Ni-NTA resin, GeneJET Plasmid Miniprep kit, GeneJET Gene extraction and DNA cleanup micro kit, and DH5 competent cells were purchased from Thermo Fisher Scientific (Waltham, MA, USA). Isopropyl β-D-1-thiogalactopyranoside (IPTG), tetracycline hydrochloride, ethylenediaminetetraacetic acid (EDTA), imidazole, Amicon Ultra centrifugal units were purchased from Merck-Millipore (Burlington, MA, USA). Dithiothreitol (DTT) was purchased from VWR (Radnor, PA, USA). Strep-Tactin®XT 4Flow high-capacity resin was purchased from IBA lifesciences (Göttingen, Germany). Restriction enzymes, T4 DNA ligase, and NEBuilder HiFi DNA assembly mix were purchased from NewEngland Biolabs (Ipswich, MA. USA).

Plasmids, pMG_cpAaLS(His12), pACYC_Ptet_cpAaLS-GFP-ssrA, pACYC_Ptet_cpAaLS-GFP, pACYC_Ptet_H-GFP-SsrA, and pACYC_Ptet_H-GFP were kindly provided by Prof. Donald Hilvert (ETH Zurich). pET15_cpAaLS(His), pMAL_H-sTEVp, TetR_Ptet_cpAaLS_MAG2, and MalE gene inserts were prepared by BioCat GmbH (Heidelberg, Germany). Other oligonucleotides were synthesized by Sigma-Aldrich (St. Louis, MO, USA) or Thermo Fisher Scientific (Waltham, MA, USA). BL21-Gold (DE3) competent cells were purchased from Agilent Technology (Santa Clara, CA, USA), and DH5 competent cells were purchased from Thermo Fisher Scientific.

An engineered human embryonic kidney (HEK) 293 cell line that possesses a firefly luciferase reporter under the control of cyclic AMP response element (CRE/CREB Luciferase Reporter HEK293 Cell Line) was purchased from BPS Biosciences (San Diego, CA, USA). Lentiviral vectors carrying a gene encoding human glucagon-like peptide-1 receptor (NM_002062) tagged with Myc-DDK peptides (GLP1R (NM_002062) Human Tagged ORF Clone) were purchased from OriGene (Rockville, MD, USA). Human GLP-1R Alexa Fluor 488-conjugated antibody was purchased from R&D Systems (Minneapolis, MN, USA).

**Molecular cloning**

Plasmid pACYC_Ptet_cpAaLS-tev-GFP-SsrA was prepared by cassette cloning via BglI and BamHI sites using pACYC_Ptet_cpAaLS-GFP-ssrA as scaffolds and oligonucleotides, FW_BglI_tev_BamHI and RV_BamHI_tev_BglI.

pET15_AaLS-wt-Strep was prepared by subcloning a gene encoding AaLS-wt-Strep from pMG_AaLS-wt-S[19] into a pET15b vector via NcoI and SpeI sites.

pACYC_Ptet_cpAaLS-tev-MAG was prepared by assembly cloning using pACYC184 as a backbone template, the insert TetR_Ptet_cpAaLS_MAG2, and oligonucleotides, FW_ACYC184 and RV_ACYC184 as primers (Supplementary Table 4).

pACYC_Ptet plasmids carrying a gene encoding cpAaLS fused to therapeutic peptides via TEV protease recognition sequence were prepared by assembly cloning using pACYC_Ptet_cpAaLS-tev*G-MAG as a backbone template, and oligonucleotides FW_XX and RV_XX (XX = LIX, TER, AVI, ENF, or THY) as insert templates, and FW_uni and RV_uni as primers (Supplementary Table 4). pET15 plasmids carrying a gene encoding cpAaLS(His) fused to therapeutic peptides via TEV protease recognition sequence were analogously prepared by assembly cloning, but using pET15_cpAaLS(His) as a backbone template FW_pET15b and RV_pET15b as primers. The inset encoding the MAG gene was prepared using pACYC_Ptet_cpAaLS-tev*G-MAG as a template and oligonucleotides, FW_MAG and RV_MAG, as primers (Supplementary Table 4). The region encoding hexahistidines in these pACYC_Ptet and pET15 plasmids, as well as pMG_cpAaLS(His12), was replaced with the one encoding strep II sequence by cassette cloning via KpnI and SacI sites using oligonucleotides, FW_Strep16 and RV_Strep16 (Supplementary Table 4). While those preliminary constructs contain additional glycine residue at the N-terminus of the peptides for efficient TEV protease cleavage, the codon for the glycine in pET15 and pACYC_Ptet was later removed by quick change PCR using oligonucleotides, FW or RV-del-peptide series (Supplementary Table 4).

pET15_H-MBP-tev*-MAG was prepared by assembly cloning using pET15_cpAaLS(His) as a backbone template, the insert MalE, and oligonucleotides, FW_pET15b_2 and RV_pET15b_2 as primers (Supplementary Table 4). The backbone of the resulting pET15_H_MBP-tev*-MAG was amplified by PCR using oligonucleotides, FW_uni_2 and RV_uni_2, and the resulting fragment was assembled with the gene fragment encoding LIX, prepared by PCR using pACYC_Ptet_cpAaLS-tev-LIX as template and oligonucleotides FW_LIX_3 and RV_LIX_3, as primers (Supplementary Table 4), yielding pET_H-MBP-tev*G-LIX. pET_H-tev*-LIX was also prepared using assembly cloning using pET15_cpAaLS(His)-tev*G-LIX as backbone template and oligonucleotides, FW_His-LIX and RV_His-LIX, as primers.

A gene encoding Strep-tagged super TEV protease variant was prepared by PCR using pMAL_H-sTEVp as template and oligonucleotides, FW_BamHI_sTEVp and RV_RV_rrnB-T1-term (Supplementary Table 2), as primers. The PCR product was subsequently subcloned into pMAL_H-sTEVp via BamHI and EcoRI sites, yielding pMAL_S-sTEVp.

The plasmids and the oligonucleotide sequences are summarized in Supplementary Tables 3 and 4. *E. coli* strain DH5α was used as the host cell for every cloning step. Sequences of plasmids were confirmed by DNA Sanger sequencing performed by Eurofins Genomics Europe Sequencing GmbH (Munich, Germany).

**Protein purification**

Cell pellet from 10-mL culture was resuspended in a 1-mL lysis buffer [50 mM Tris-HCl buffer (pH 8.0) containing 300 mM NaCl and 1 mM EDTA], supplemented with 0.1 mg/mL lysozyme and 0.01 mg/mL DNase I. After 1h at room temperature, the cells were lysed by sonication with a Sonics & Materials VCX 130 ultrasonic processor (Newtown, CT, USA) using a 5/1-sec on/off cycle and a 40% amplitude on ice for 2 min. After removal of the insoluble fraction by centrifugation at 21,500 ×g and 25 °C for 10 min, the supernatant was loaded on a StrepTactin XT resin (0.2 mL) pre-equilibrated with the lysis buffer in a gravity column. The protein-bound resin was washed with the lysis buffer (5-column volume), and the Strep-tagged AaLS was eluted with the lysis buffer supplemented with 50 mM D-biotin. The sample buffer was subsequently replaced with 50 mM Tris-HCl buffer (pH 8.0) containing 0.3 mM NaCl and 5 mM EDTA using Amicon Ultra-0.5 centrifugal unit (50 k MWCO). The resulting solution was used as a stock for further analysis by SDS-PAGE, native agarose electrophoresis (AGE), absorbance spectrometry, and transmission electron microscopy (TEM). AaLS concentration was determined by absorbance at 280 nm using the extension coefficients listed in Supplementary Table 1.

Purification of cpAaLS(His) was performed using the protocol described previously[20].

For guest retrieval experiments, protein purification using affinity chromatography was performed using an analogous protocol starting with cell pellets from 200-mL or 500-mL cultures for GFP or LIX, respectively. After elution, the buffer was replaced with 50 mM Tris-HCl buffer (pH 8.0) containing 0.3 mM NaCl, 1 mM EDTA, and 2 mM DTT for the following TEV protease treatment.

**GFP loading efficiency calculation[27]**

The number of GFP encapsulated in an AaLS cage is defined in the following equation.

$$\#\text{GFP per AaLS assembly} = [\text{GFP}] / ([\text{AaLS}] / n)$$

where $n$ indicates the number of building blocks to form an AaLS assembly, e.g. $n = 60$ for the AaLS-wt dodecahedron. Data shown in Figure 2f are provided as the number of GFP per AaLS pentamer ($n = 5$) to disregard the differences in the number of protomers to form a cage-like assembly between AaLS-wt (60) and cpAaLS(His) (120 or 180). Following the Lambert-Beer's law, GFP and AaLS concentrations are given by the following equations.

$$[\text{GFP}] = (A_{475\text{obs}} / \varepsilon_{\text{G475}} \times l)$$

$$[\text{AaLS}] = (A_{280\text{obs}} - \varepsilon_{\text{G280}} \times [\text{GFP}] \times l) / \varepsilon_{\text{A280}} \times l$$

where $A_{475obs}$ and $A_{280obs}$ are the observed absorbance values at 475 nm and 280 nm, respectively; $l$ is light pass (= 1 cm); $\varepsilon_{G475}$ and $\varepsilon_{G280}$ are the molar extinction coefficients of GFP at 475 nm and 280 nm, respectively; $\varepsilon_{A280}$ is the extinction coefficient of AaLS (Supplementary Table 1). The reported extinction coefficients of the red-shifted GFP, $\varepsilon_{G280}$ = 30,600 $M^{-1}cm^{-1}$ and $\varepsilon_{G475}$ = 56,600 $M^{-1}cm^{-1}$, were used for the calculation[20].

**LIX purification and characterization**

The cpAaLS(Strep)-LIX sample in 50 mM Tris-HCl (pH 8.5) containing 1 mM EDTA and 2 mM DTT was added with super TEV protease possessing strep-tag (1:15 as mass ratio) and kept at room temperature overnight. After the addition of 10% volume of ethanol, the insoluble fraction was removed by centrifugation at 25 °C and 16,000 ×g for 10 min. The supernatant was passed through StrepTactin XT resin, followed by lyophilization. The crude peptide was then dissolved in 20% MeCN aqueous solution supplemented with 1% (v/v) trifluoroacetic acid (TFA), and purified by reverse-phase high-performance liquid chromatography (RP-HPLC) [column: Nucleosil C8, 120 Å, 5 µm, 4.6 × 150 mm (Macherey Nagel, Düren, Germany); flow rate: 1 mL/min; eluent A: $H_2O$ containing 0.1% TFA; eluent B: MeCN containing 0.08% TFA; gradient: 25-50 %B over 25 min]. The purity of the peptide was confirmed by RP-HPLC [column: Nucleosil C18, 120 Å, 5 µm, 4.6 × 150 mm (Macherey Nagel); flow rate: 1 mL/min; eluent A: $H_2O$ containing 0.1% TFA; eluent B: MeCN containing 0.08% TFA; gradient: 5-95 %B over 30 min]. The identity of the purified LIX was confirmed by electron spray ionization time-of-flight mass spectrometry (ESI-TOF-MS) on a Bruker Daltonics micrOTOF-Q II: 4856.6418 [calcd 4856.5283].

**HEK cell culture**

The CRE/CREB luciferase reporter HEK293 cell line was routinely maintained in Dulbecco's modified Eagle's medium with high glucose (DMEM-HG) supplemented with 10% heat-inactivated fetal bovine serum (FBS), 100 U/mL penicillin and 0.1 mg/mL streptomycin, referred to as complete medium (CM) hereinafter, at 37°C in a humidified 5% $CO_2$ atmosphere.

**Transduction of the CRE/CREB Reporter HEK293 cell lines with GLP-1R-lentivirus**

CRE/CREB luciferase reporter HEK cells were seeded on a 12-well plate (250,000 cells per well) and cultured in CM at 37°C overnight. The next day, the medium was exchanged with CM (without antibiotics) supplemented with lentiviral particles (~$10^6$ transduction units) and 5 mg/mL polybrene, and the cells were incubated for 2 days. The transduction medium was then replaced with CM containing 2 μg/mL puromycin and cultured overnight. The cells were then transferred to a 6-well plate and cultured in CM containing 2 μg/mL puromycin for an additional 3 days. The resulting cells were added with 10% dimethyl sulfoxide and stored in liquid nitrogen.

**GLP-1R expression test**

The lentivirally transduced CRE/CREB luciferase reporter HEK cells (approximately $5 \times 10^5$ cells) in 100 μL Dulbecco's modified phosphate-buffered saline (DPBS) were treated with Alexa488-conjugated anti-GLP-1R antibody (2 μL, 1:50 dilution) at room temperature for 30 min. The cells were washed thrice with DPBS and resuspended in 2 mL DPBS. Propidium iodide was added to the cell suspension to the final 1 μg/mL to discriminate the dead cell population during the analysis. Flow cytometry of the cells was performed on a Navious Flow Cytometer (Beckman Coulter, Brea, CA, USA). A blue laser operating at 488 nm was used for excitation, and fluorescence was detected through a 525/40 nm emission filter. Data were analyzed using Kaluza C software (Beckman Coulter), and the results from $8.3 – 10.0 \times 10^3$ cells are provided in Supplementary Figure 11.

#### Uncropped gel images

##### Uncropped gel images 1 (Figure 2e)

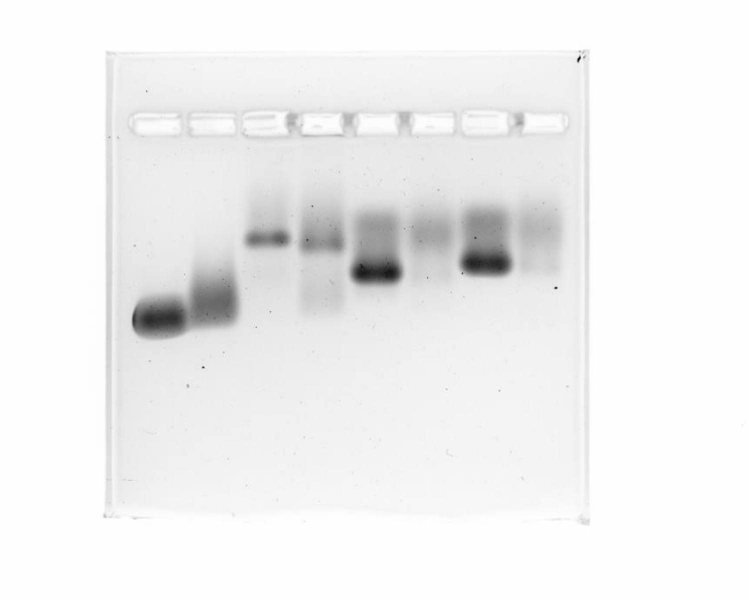

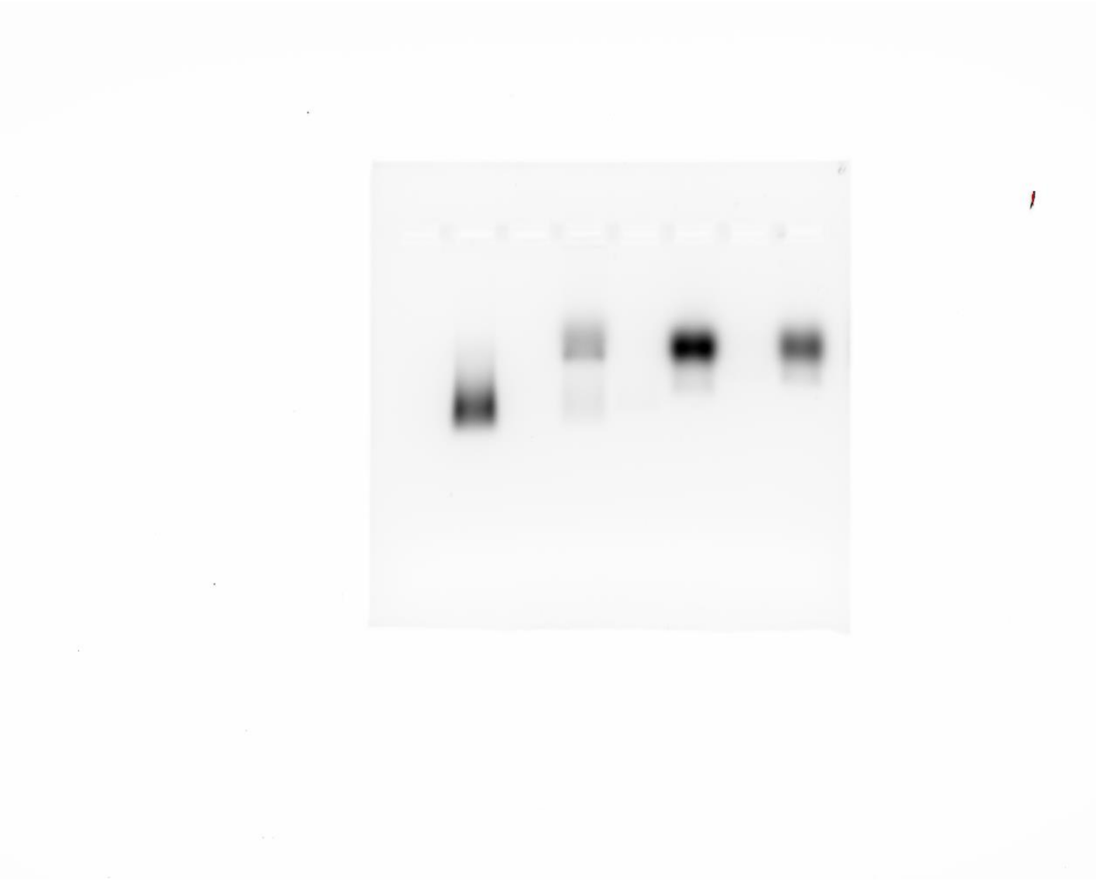

**Uncropped gel images 2 (Figure 3c)**

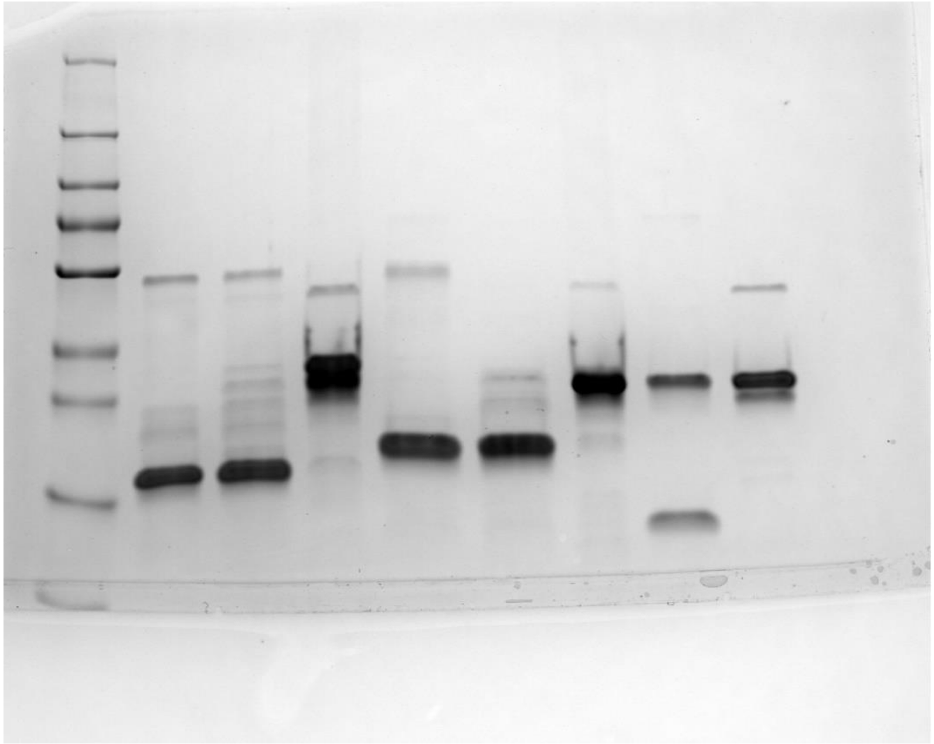

**Uncropped gel image 3 (Figure 4b and Supplementary Figure 6)**

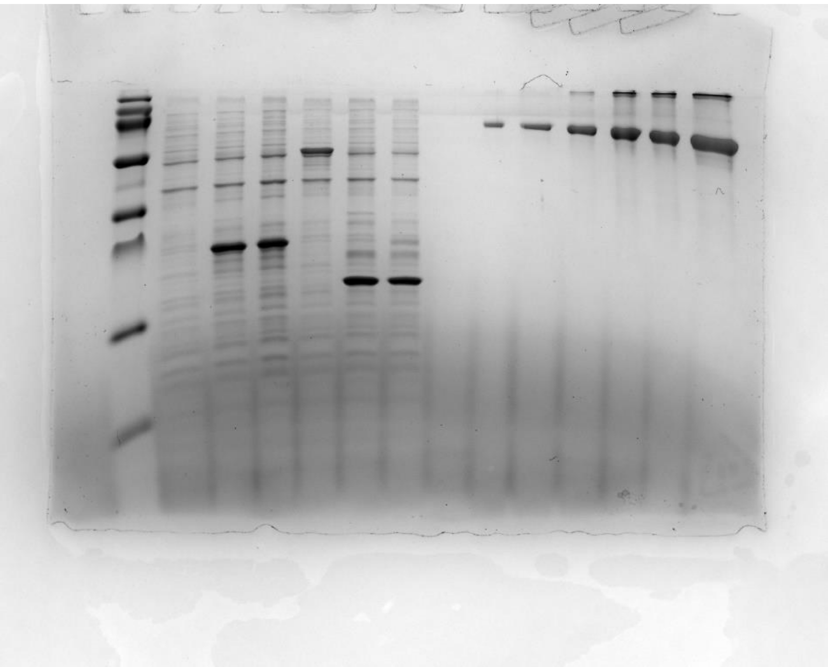

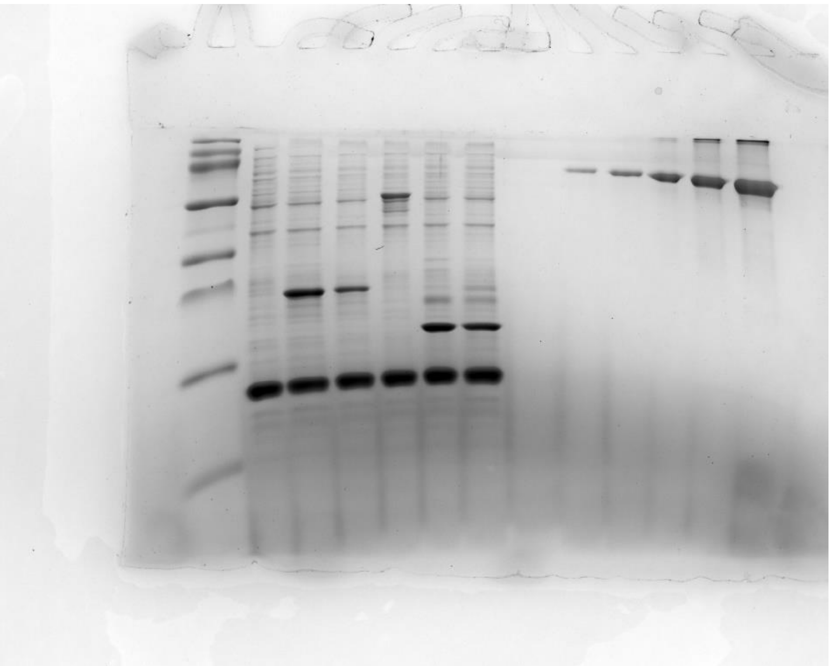

**Uncropped gel image 4 (Figure 5a and Supplementary Figure 7a)**

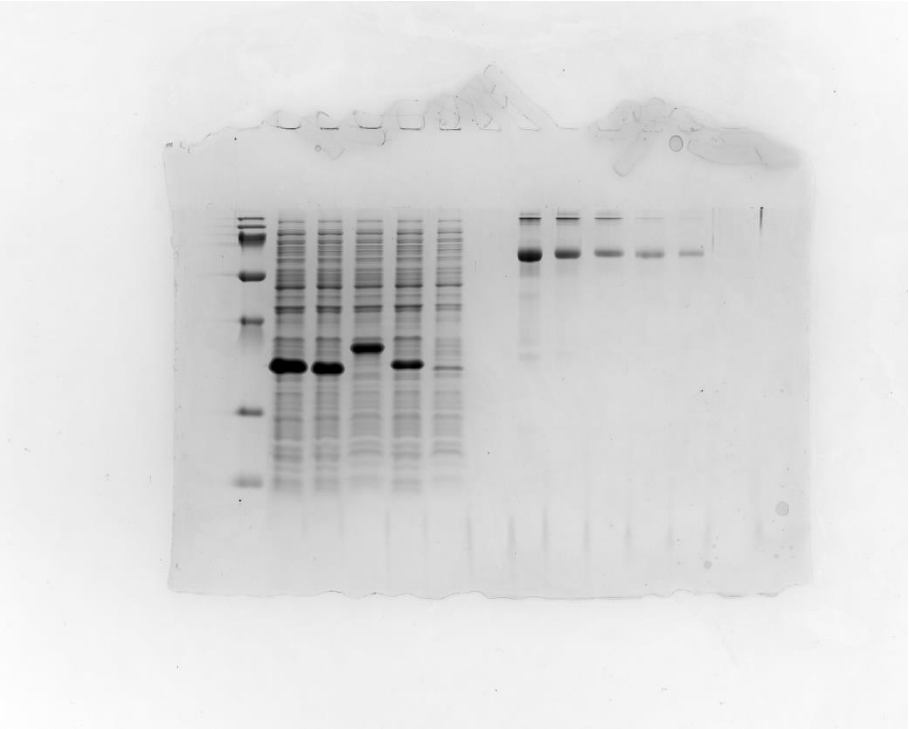

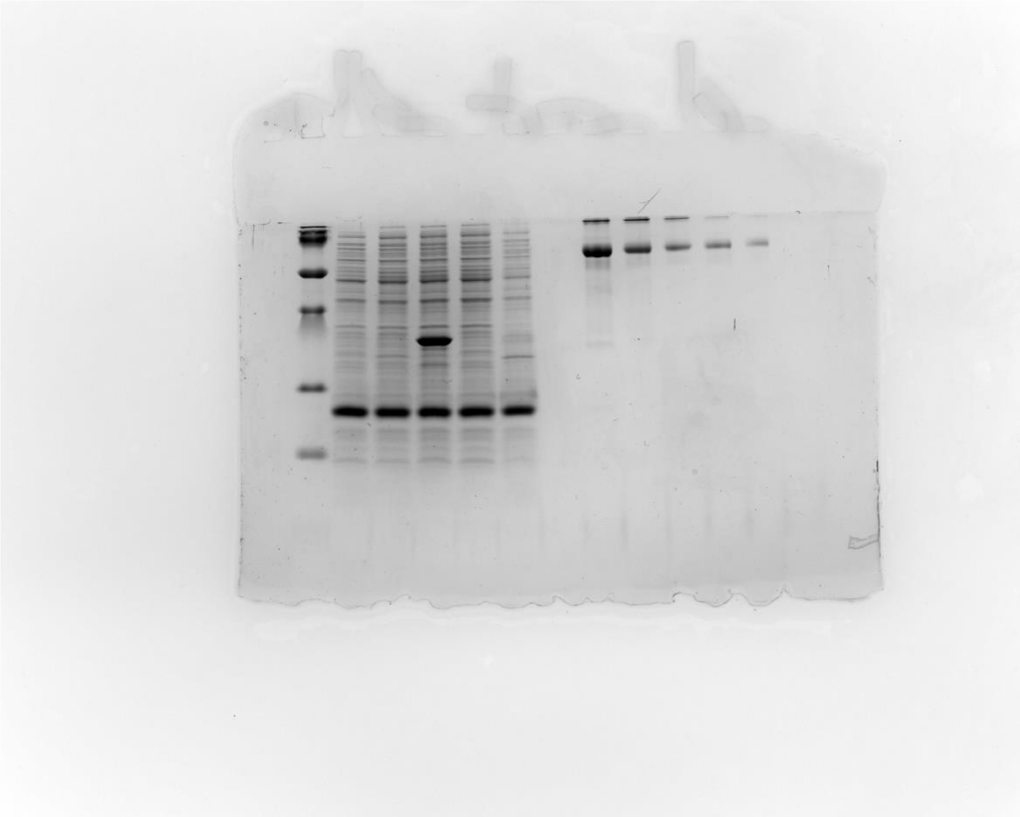

**Uncropped gel image 5 (Figure 5c and Supplementary Figure 8a,b, left gels)**

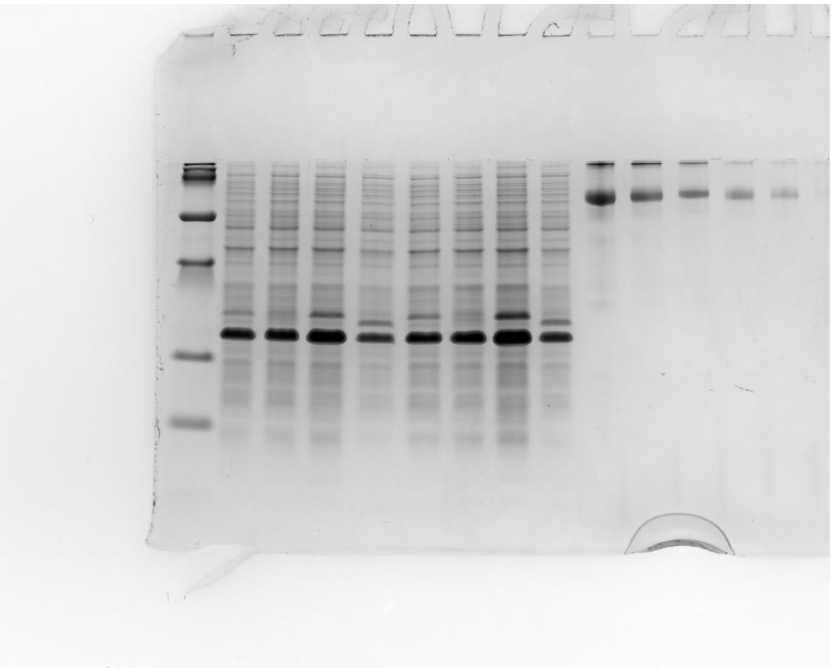

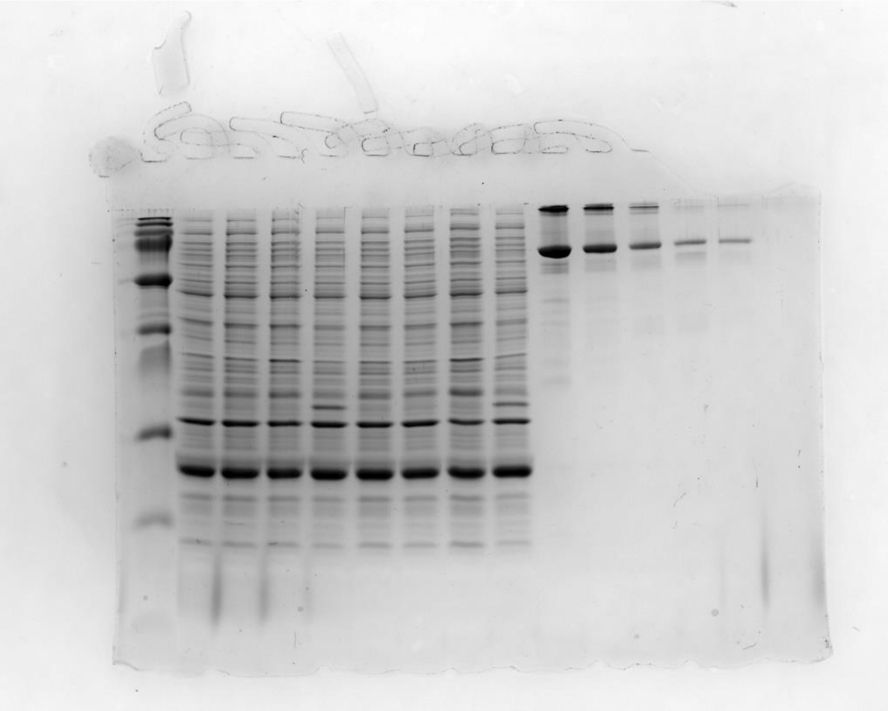

**Uncropped gel image 6 (Supplementary Figure 8a,b, right gels)**

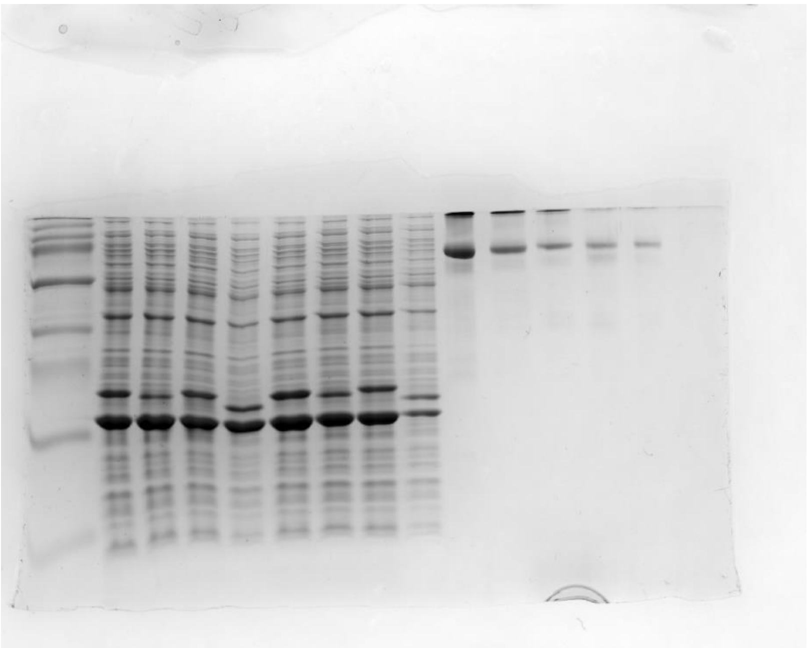

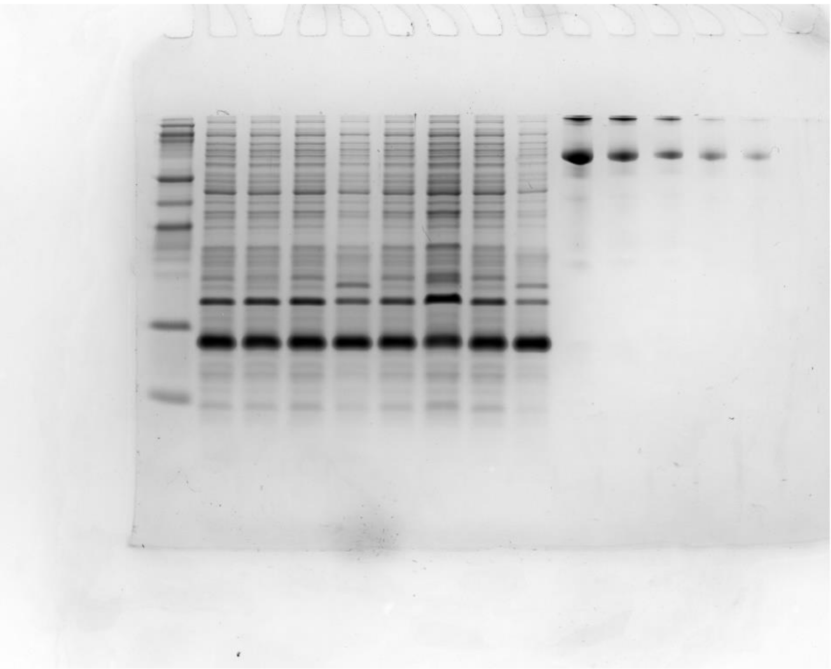

**Uncropped gel image 7 (Figure 6a and Supplementary Figure 7a)**

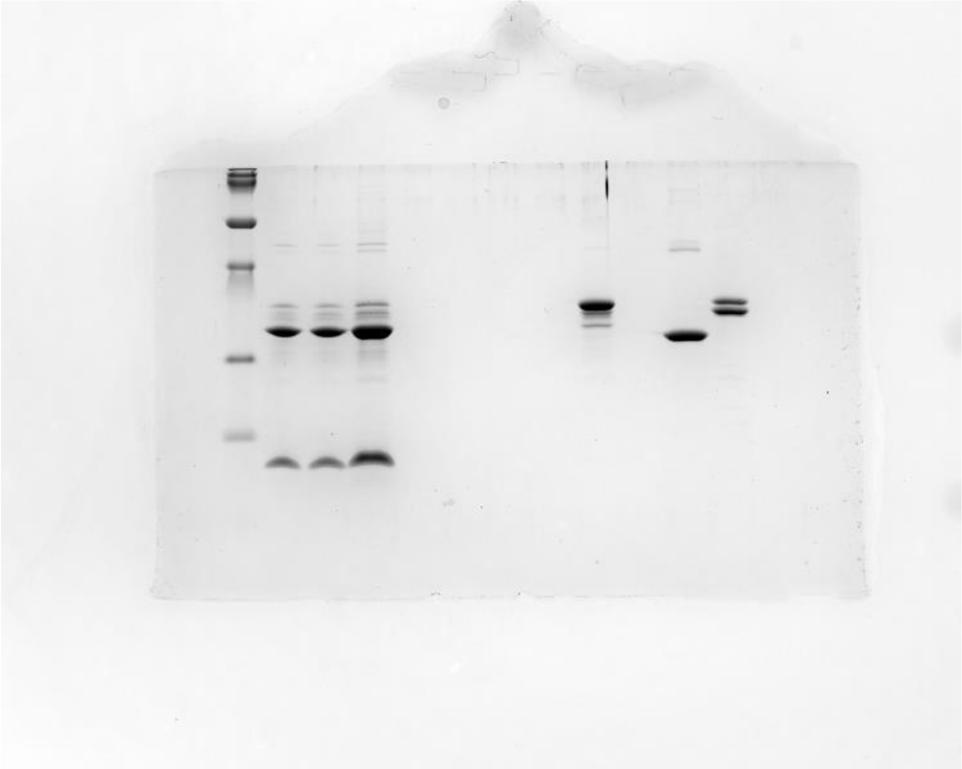

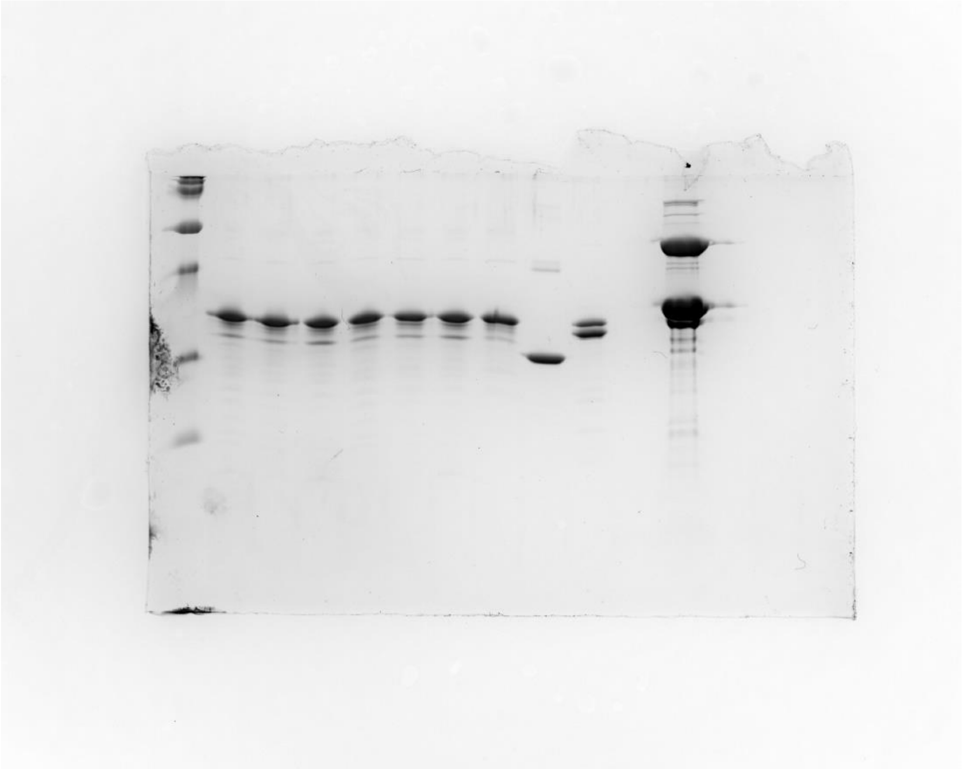
